# Compartment-specific soluble immune profiles associated with preterm birth, perinatal death, and low birthweight in pregnant individuals living with HIV

**DOI:** 10.64898/2026.05.04.722284

**Authors:** Jacqueline Corry, Natalia Zotova, Martine Tabala, Fidéle Lumande Kasindi, Bienvenu Lebwaze Massamba, Pelagie Babakazo, Jennifer A. Manuzak, Namal P.M. Liyanage, Nicholas T. Funderburg, Marcel Yotebieng, Jesse J. Kwiek

## Abstract

**Background:** Human immunodeficiency virus (HIV) infection in pregnancy is associated with preterm birth (PTB), low birthweight (LBW), and perinatal death (PND). Although antiretroviral therapy (ART) suppresses viral load it does not prevent HIV-associated adverse pregnancy outcomes or resolve inflammation. As circulating maternal immune factors may not fully capture maternal-fetal interface immune dysregulation, this observational cohort study aimed to identify localized and systemic immune factors associated with PTB, LBW and PND in ART-treated pregnant people living with HIV (PPLWH).

**Methods:** We enrolled 118 PPLWH in Kinshasa, Democratic Republic of the Congo, during the second or third trimester. We collected maternal peripheral plasma (at enrollment, 1-3 days post-delivery, and postpartum) alongside umbilical cord and placental plasma at delivery. Concentrations of 45 immune factors were measured via LegendPlex and ELISAs and associations analyzed using Kruskal-Wallis tests with Dunn’s correction or Mann-Whitney tests.

**Results:** Placental plasma exhibited the highest overall concentrations of immune factors, highlighting a distinct localized microenvironment. Among 118 pregnancies, 35 (30%) resulted in PTB, 10 (9%) in PND, and 9 (8%) in LBW. Compared to term births, PTB was associated with higher levels of the chemokines CCL20, CXCL9, and CXCL10 in cord and/or postdelivery plasma (p<0.01), while placental CCL20 levels were lower (p<0.05). Compared to live births, PND was associated with higher postdelivery CXCL1, cord IL-8, placental MPO and NGAL (p<0.05); higher postdelivery CXCL5 (p<0.01); and higher S100A8/A9 levels in cord and postdelivery plasma (p<0.01 and p<0.001, respectively). Finally, LBW was associated with higher enrollment IL-18 and S100A8/A9 levels (p<0.05 and p<0.01, respectively); as well as higher SAA levels in postdelivery and postpartum plasma (p<0.05).

**Conclusions:** In ART-treated PPLWH, distinct adverse birth outcomes are driven by time- and compartment-specific immune pathways. PTB is associated with localized T-cell chemokine responses, PND with neutrophil recruitment and activation, and LBW with pro-inflammatory cytokine and acute-phase protein responses. These pathways provide mechanistic insights into pregnancy complications in PPLWH and highlight potential compartment-specific biomarkers for risk stratification.

## Background

Women and girls accounted for 53% of the roughly 40.8 million people living with HIV globally at the end of 2024 [1]. The health impact on this demographic is particularly severe in sub-Saharan Africa, a region which not only accounted for about 80% of all new infections among young women and girls in 2024 [1], but which is also concurrently struggling with a disproportionate malaria burden [2]. In non-pregnant individuals, HIV affects the immune system in a variety of ways, eventually leading to chronic inflammation [3–6]. Pregnancy, however, requires precise immune regulation to maintain the delicate balance necessary to sustain the fetal allograft [7–10]. HIV disrupts this immune balance [11–15], which can lead to poor placental development and abnormalities [16–18], all of which have been associated with adverse pregnancy outcomes [17,19–26], including vertical HIV transmission, preterm birth (PTB), low birth weight (LBW), stillbirth, and maternal, neonatal, and infant mortality [27–38]. The risk of poor fetal outcomes is further compounded in pregnant individuals coinfected with the malaria-causing parasite *Plasmodium*, as these parasites can accumulate within the placenta [39–43]. Although antiretroviral therapy (ART) effectively decreases HIV viral load, and minimizes vertical transmission, ART is less effective at mitigating the chronic inflammation associated with HIV infection [4–6,44–46] and preventing HIV-associated adverse pregnancy outcomes [30,37,47–54].

Adverse pregnancy outcomes carry significant economic, health, and social costs. In 2021, PTB complications continued to be a leading cause of death in both lower- and lower-middle-income countries [55]. Not only have both PTB and LBW been associated with an increased infant mortality risk [31,56], but both are also associated with long-term health complications that can persist into childhood and adulthood [31,57,58]. These impacts are not limited to the child; individuals who experience fetal demise or neonatal death face a higher risk of perinatal loss in subsequent pregnancies [59,60]. Furthermore, emerging evidence in population-based cohorts suggests that adverse pregnancy outcomes may be associated with early-onset cardiovascular disease or death in the gestational parent [61].

Despite the severity of these complications, previous studies characterizing adverse birth outcomes in PPLWH have primarily focused on a single outcome of interest, measured only a small number of immune factors, or failed to evaluate plasma across multiple critical time points and anatomical compartments. To address these gaps, here we comprehensively tested the concentrations of 45 immune factors in the blood of PPLWH on ART across multiple stages: during pregnancy, postdelivery, and postpartum. We also assessed these factors in umbilical cord blood, which represents the fetal compartment, and placental blood, which represents primarily maternal blood enriched with placenta-secreted factors. With these distinct plasma samples, we sought to determine how immune factor concentrations differ between term and PTB, live births and perinatal death (PND), and normal birthweight and LBW.

## Methods

### Definitions

For clarity, we categorized key concepts into immunological, obstetrical, and infection-related definitions. Immunologically, cytokines are defined as peptides or proteins secreted by cells that can have autocrine, paracrine or endocrine effects on target cells that express the requisite receptor(s). Pro-inflammatory cytokines induce, or trigger inflammation, whereas anti-inflammatory cytokines decrease it. Chemokines are a subset of cytokines that induce directional movement along a gradient. Damage-associated molecular patterns (DAMPs) are intracellular molecules that are released from dead or dying cells; they trigger inflammation by binding to pattern recognition receptors to signal cellular stress or tissue injury [62]. Acute phase proteins (APPs) are a class of proteins whose concentration in blood increases in response to injury, trauma, or infection [63]. In this manuscript, “biomarkers” refers to non-cytokine molecules whose measurable blood concentrations change in response to inflammation but which neither bind cytokine receptors nor actively drive the inflammatory process. The term “immune factor” is the collective name for all analyzed cytokines, chemokines and biomarkers.

In terms of obstetrical definitions, gestational age was based on the self-reported last menstrual period at enrollment when available (n=95), or clinical estimates when not (n=23). Births prior to 37 weeks’ gestation were classified as preterm births (PTBs); those at or after 37 weeks as term birth. Perinatal death (PND) encompassed stillbirth (fetal death after 28 weeks but prior to or during birth [64]) and neonatal death (death within the first 28 days of birth). Live birth includes all neonates that survived 48 h after birth. Neonate weights at delivery were categorized as low birth weight (LBW, defined as ≤2500g), normal birth weight (defined as >2500g), or missing for neonates lacking birthweight information. Blood draws occurred at enrollment (between gestational weeks 15 and 40), 1–3 days postdelivery (PD), and approximately 6 weeks postpartum (median 6.7 weeks; 95% CI 6-10 weeks). Cord blood refers to blood collected from the umbilical cord that represents the fetal compartment, collected at time of delivery, and placental blood—primarily maternal blood from the intervillous space—was collected immediately after the cord blood [65].

The technical limit of detection for HIV RNA was <40 HIV RNA copies/ml of blood, (see below for methods). Placental *Plasmodium* infection classifications were made by a pathologist and were based on Bulmer’s classification [66]: an *active infection* was characterized by the presence of parasites in maternal erythrocytes within the intervillous spaces and malarial pigment (hemozoin) in circulating erythrocytes and/or monocytes, with a notable absence of hemozoin or plasmodial cells in fibrin. An *active-chronic infection* exhibited the same characteristics as an active infection but additionally featured the presence of hemozoin or plasmodial cells in fibrin, the syncytiotrophoblasts, and/or the stroma. Finally, a *past-chronic infection* was characterized by a lack of parasites in erythrocytes alongside the presence of hemozoin pigment or plasmodial cells confined to fibrin.

### Study setting

This study was an observational cohort study nested within the larger randomized control trial for the Continuous Quality Improvement-Prevention of Mother to Child Transmission (CQI-PMCT) [67–73]. The parent study was an open label, parallel, group randomized trial designed to evaluate continuous quality improvements (CQI) on long-term ART outcomes among pregnant and breastfeeding individuals receiving care in Kinshasa province.

### Study Design, Participants, and Specimen Collection

Drawing from the CQI-PMCT framework, we identified all pregnant individuals in their second or third trimester living with HIV and receiving care at any of the participating maternal and child health facilities between October 2020 and May 2021. After obtaining written consent, 199 eligible participants were enrolled. Of these, 81 were excluded from the final analysis due to the availability of only a single plasma sample (n=28), missing important delivery information (e.g., birth weight, baby’s sex, mode of delivery, or pathologic information on the placenta/umbilical cord; n=48), or multiple births (n=5). The final cohort for this analysis consisted of 118 participants.

To monitor these participants longitudinally, trained personnel administered questionnaires and collected maternal peripheral venous blood and dried blood spots at three specific time points: enrollment, 1–3 days postdelivery, and approximately 6 weeks postpartum. Venous blood was drawn into EDTA-containing tubes, centrifuged for 10 minutes at 200 × g, and the resulting plasma was aliquoted and stored at -80°C. Concurrently, DBS were obtained via finger prick on Whatman 903 protein saver cards (Cytiva), dried at room temperature for at least three hours, then packaged with desiccant and stored at -20°C. Postdelivery dried blood spots were tested for HIV viral RNA using the m2000rt Real-Time HIV-1 assay (Abbott) by the National AIDS Reference Laboratory in Kinshasa.

At delivery the umbilical cord and placenta were collected. The cord was washed with normal sterile saline and approximately 3 cm from its insertion into the placenta a 5 ml syringe was inserted into the umbilical vein with a 20-22 gauge needle was used to aspirate cord blood into an EDTA collection tube. Following cord blood collection, the placental prick method described by Othoro *et al.* was employed to collect blood from the intervillous space into EDTA-containing tubes [65]. Cord and placental blood were centrifuged for 10 mins at 200 x g, and plasma was aliquoted and stored at -80°C. Placentas were weighed and 2 x 1 x 0.5 cm sections were taken from the maternal and fetal surfaces as previously described [74]. The Bulmer histological classification [66] was used to classify placental plasmodial lesions.

### Specimen processing and laboratory testing

Following sample collection in the DRC, all plasma aliquots were shipped on dry ice to The Ohio State University (Columbus, Ohio) and stored at -80°C. Prior to assaying, samples were thawed and centrifuged twice at 2,500 x g for 5 minutes to remove particulates.

Immune factor names and abbreviations were standardized via UniProt (*Homo sapiens*) following the International Protein Nomenclature Guidelines [75]; full details including units of measure and kit purchasing information are provided in **Table S1**. The LEGENDplex Human Proinflammatory Chemokine Panel 1 (BioLegend, 740984), LEGENDplex COVID-19 Cytokine Storm Panel 1 & 2 (BioLegend, 741095) and LEGENDplex Vascular Inflammation panel 1 (BioLegend, 740551) were used to measure immune factors according to the manufacturer’s filter plate protocol as previously described [76]. Briefly, data were collected on the MACS Quant 10 Flow cytometer (Miltenyi Biotec) and using MacsQuant Software (Miltenyi Biotec). Analysis was performed using LEGENDplex online Qognit software (BioLegend, Qognit Inc.) based on standards; gates were then applied to all standards and samples and five-parameter logistic standard curves were generated for all and concentrations were interpolated from these curves. Additionally, seven specific immune factors (CXCL13, IL-4, IFN-λ-1, sCD14, sCD163, sTNFRSF1A, and sTNFRSF1B) were individually measured via DuoSet ELISA (R&D Systems) with minor modifications [76]. Absorbances were read on a SpectraMax i3x (Molecular Devices). See **Table S2** for dilution factors, limits of detection (LoD), sample number, and samples (n and %) above and below the LoD.

We calculated a Composite Inflammatory Score to quantify overall inflammation. For the 26 pro-inflammatory factors (detailed in **Figure 1** and **Table S1**), individuals received a score of +1 if their concentration was above the 75th percentile, and -1 if it was below the 25th percentile. Conversely, for the three anti-inflammatory factors, a score of -1 was assigned for concentrations above the 75th percentile, and +1 for those below the 25th percentile. The remaining 16 factors, categorized as mixed pro-/anti-inflammatory or general biomarkers, were not scored (**Table S9**). Total scores were calculated for each individual across each sample type. Participants were categorized by adverse birth outcomes; these groups were not mutually exclusive, meaning individuals could appear in multiple categories. Total median scores were calculated per outcome group (excluding participants missing birthweight data with no other adverse outcomes) and visualized using a stacked bar graph.

**Figure 1.**
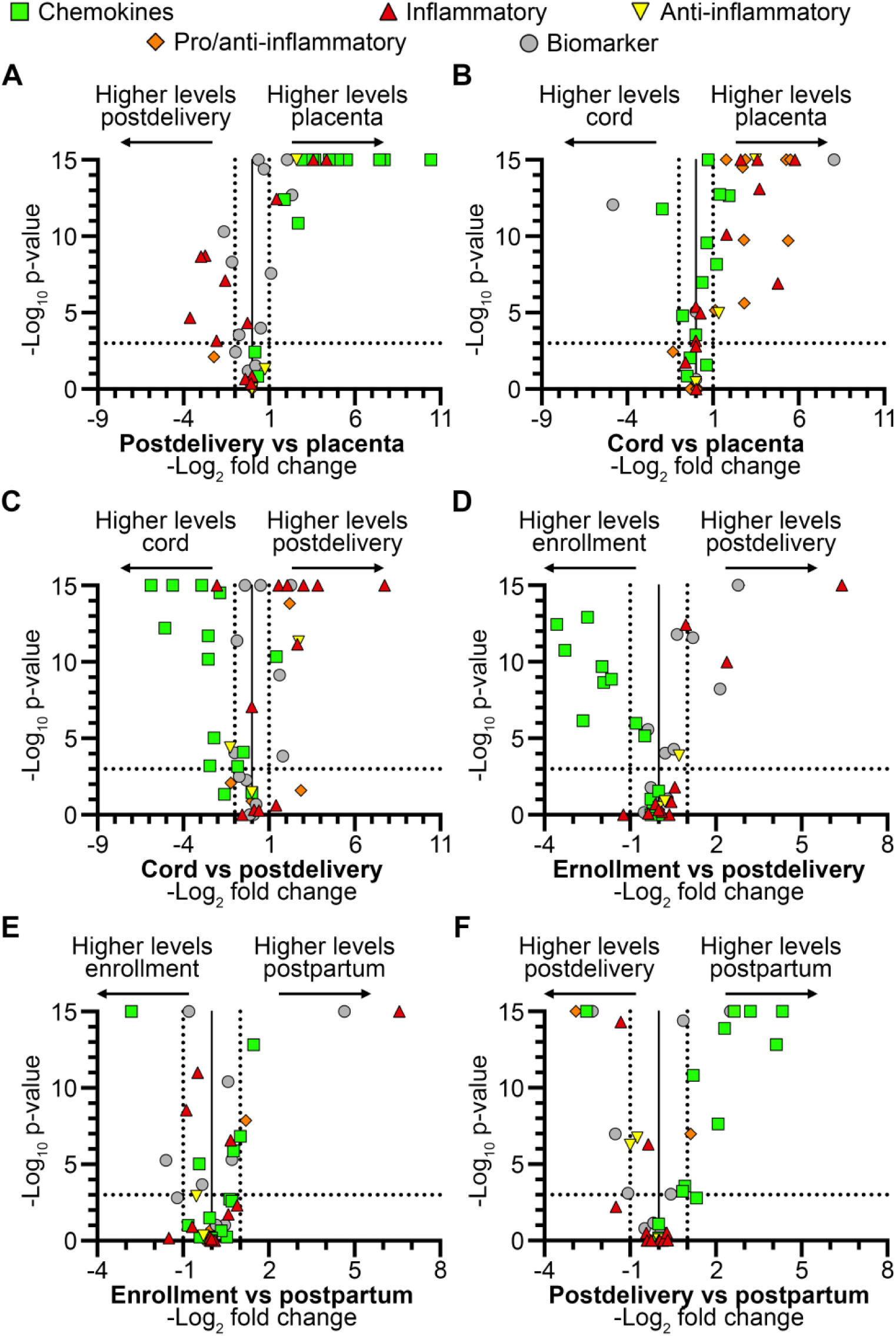
Volcano plots showing differential expression of immune factors in plasma from different locations. Differential expression plots displaying the -log_10_ of the p-value of and -log_2_ magnitude fold change in immune factors between (A) postdelivery and placental plasma, (B) cord and placental plasma, and (C) cord and postdelivery plasma, (D) enrollment and postdelivery plasma, (E) enrollment and postpartum plasma, or (F) postdelivery and postpartum plasma. Statistical significance was tested by Kruskal-Wallis with a Dunn’s correction for multiple comparison. Vertical dotted lines are equivalent to a 2-fold change, horizontal dotted lines are set to a p-value that is equivalent to 0.001. Immune factor groups are defined in **Table S1**.

### Statistical Analyses

Data processing was performed in Microsoft Excel, while statistical testing and visualization were conducted in GraphPad Prism (v. 10.4.1–10.6.0). Descriptive statistics for immune factor data, adverse birth outcomes and Composite Inflammatory Score are available in **Tables S3** and **S6-S9**. Depending on the data type and distribution, significance was evaluated using two-tailed Mann-Whitney tests, two-sided Fisher’s exact tests, Kruskal-Wallis tests with or without Dunn’s correction for multiple comparisons, or two-tailed Spearman correlations (significance cutoffs: rs < -0.2 or > 0.2, p < 0.05). Because this is an exploratory study, we did not correct for multiple hypothesis testing across all findings [77].

## Results

### Participant characteristics

We recruited 118 PPLWH during their second or third trimester. At delivery, 35 neonates were delivered prematurely, including one stillborn fetus, one neonatal death, one LBW neonate, and five neonates missing birth weight data. Across the entire cohort, there were ten PNDs and nine LBW neonates. Only 25 PPLWH had a detectable HIV viral load at delivery, and of those, 10 had a viral load below 1000 HIV RNA copies/mL. Additionally, most participants showed evidence of past or present placental malaria infection (**Table 1**). Term and PTB cases varied by both date of enrollment and birthweight. Finally, the mode of delivery differed between PND cases and live births (**Table 2**).

**Table 1.**
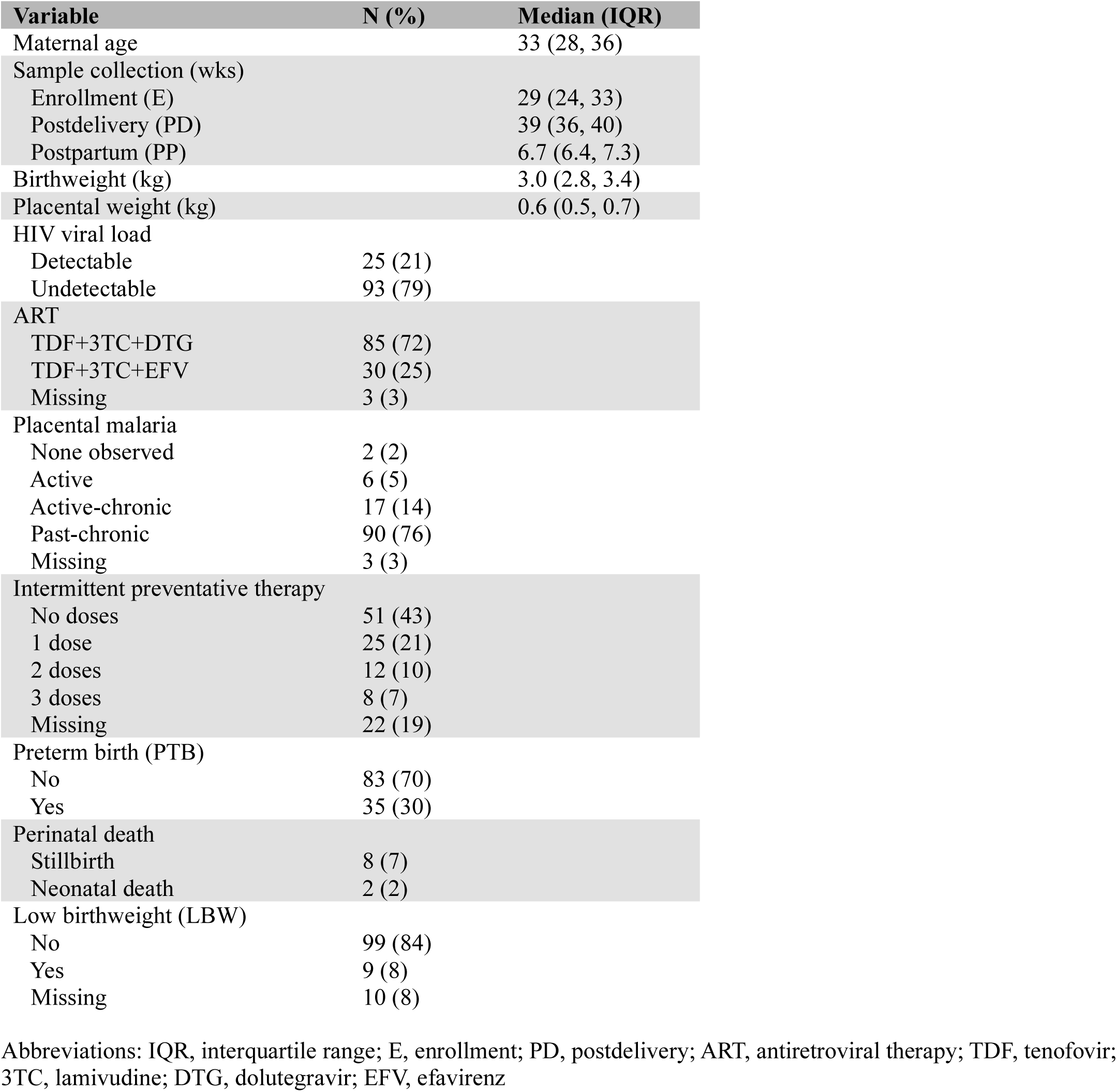
Obstetrical data and outcome.

| Variable | N (%) | Median (IQR) |
| --- | --- | --- |
| Maternal age |  | 33 (28, 36) |
| Sample collection (wks) |  |  |
| Enrollment (E) |  | 29 (24, 33) |
| Postdelivery (PD) |  | 39 (36, 40) |
| Postpartum (PP) |  | 6.7 (6.4, 7.3) |
| Birthweight (kg) |  | 3.0 (2.8, 3.4) |
| Placental weight (kg) |  | 0.6 (0.5, 0.7) |
| HIV viral load |  |  |
| Detectable | 25 (21) |  |
| Undetectable | 93 (79) |  |
| ART |  |  |
| TDF+3TC+DTG | 85 (72) |  |
| TDF+3TC+EFV | 30 (25) |  |
| Missing | 3 (3) |  |
| Placental malaria |  |  |
| None observed | 2 (2) |  |
| Active | 6 (5) |  |
| Active-chronic | 17 (14) |  |
| Past-chronic | 90 (76) |  |
| Missing | 3 (3) |  |
| Intermittent preventative therapy |  |  |
| No doses | 51 (43) |  |
| 1 dose | 25 (21) |  |
| 2 doses | 12 (10) |  |
| 3 doses | 8 (7) |  |
| Missing | 22 (19) |  |
| Preterm birth (PTB) |  |  |
| No | 83 (70) |  |
| Yes | 35 (30) |  |
| Perinatal death |  |  |
| Stillbirth | 8 (7) |  |
| Neonatal death | 2 (2) |  |
| Low birthweight (LBW) |  |  |
| No | 99 (84) |  |
| Yes | 9 (8) |  |
| Missing | 10 (8) |  |

Abbreviations: IQR, interquartile range; E, enrollment; PD, postdelivery; ART, antiretroviral therapy; TDF, tenofovir; 3TC, lamivudine; DTG, dolutegravir; EFV, efavirenz

**Table 2.** Obstetrical data by adverse outcome☨.

| Variable | T (n=83) | PTB (n=35) | p-value<br>‡ | LB (n=108) | PND (n=10) | p-value<br>‡ | N (n=99) | LBW (n=9) | M (n=10) | p-value<br>α |
| --- | --- | --- | --- | --- | --- | --- | --- | --- | --- | --- |
| Maternal age | 32 (28, 36) | 34 (28, 37) | 0.24 | 33 (27, 36) | 34 (32, 39) | 0.39 | 33 (28, 37) | 34 (26, 35) | 34 (27, 37) | 0.97 |
| Sample collection |  |  |  |  |  |  |  |  |  |  |
| Enrollment (E) | 30 (25, 36) | 26 (22, 32) | **0.003** | 29 (24, 33) | 33 (27, 37) | 0.1 | 29 (24, 34) | 31 (29, 35) | 25 (21, 26) | 0.05 |
| Postdelivery (PD) | 39 (38, 41) | 35 (33, 36) | **<0.0001** | 39 (36, 40) | 38 (37, 42) | 0.64 | 39 (36, 40) | 39 (39, 40) | 37 (35, 38) | 0.13 |
| Postpartum (PP) | 6.7 (6.4, 7.3) | 6.7 (6.4, 7) | 0.73 | 6.7 (6.4, 7.3) | 6.6 (6.6, 7.3) | 0.8 | 6.7 (6.4, 7.3) | 6.9 (6.5, 7) | 6.4 (6.4, 7.4) | 0.96 |
| Birth weight (kg) | 3.1 (3, 3.4) | 2.9 (2.8, 3.2) | **0.02** | 3.1 (2.8, 3.4) | 3.1 (2.9, 3.3) | 0.94 | 3.1 (3, 3.4) | 2.5 (2.4, 2.5) | N/A | **<0.0001** |
| Placental weight (kg) | 0.6 (0.5, 0.7) | 0.5 (0.5, 0.7) | 0.14 | 0.6 (0.5, 0.7) | 0.6 (0.5, 0.7) | 0.68 | 0.6 (0.5, 0.7) | 0.5 (0.4, 0.5) | 0.6 (0.5, 0.7) | 0.08 |
| Mode of delivery |  |  |  |  |  |  |  |  |  |  |
| Cesarian | 13 (16) | 9 (26) | 0.55 | 19 (18) | 3 (30) | **0.003** | 21 (21) | 0 (0) | 1 (10) | 0.62 |
| Dystocia | 2 (2) | 0 (0) |  | 0 (0) | 2 (20) |  | 2 (2) | 0 (0) | 0 (0) |  |
| Eutocia | 67 (81) | 26 (74) |  | 88 (81) | 5 (50) |  | 75 (76) | 9 (100) | 9 (90) |  |
| Missing | 1 (1) | 0 (0) |  | 1 (1) | 0 (0) |  | 1 (1) | 0 (0) | 0 (0) |  |
| Gravidity |  |  |  |  |  |  |  |  |  |  |
| Primigravid | 17 (20) | 7 (20) | >0.99 | 23 (21) | 1 (10) | 0.68 | 21 (21) | 2 (22) | 1 (10) | 0.81 |
| Multigravid | 66 (80) | 28 (80) |  | 85 (79) | 9 (90) |  | 78 (79) | 7 (78) | 9 (90) |  |
| HIV-viral load |  |  |  |  |  |  |  |  |  |  |
| Detectable | 16 (19) | 9 (26) | 0.47 | 23 (21) | 2 (20) | >0.99 | 20 (20) | 2 (22) | 3 (30) | 0.73 |
| Undetectable | 67 (81) | 26 (74) |  | 85 (79) | 8 (80) |  | 79 (80) | 7 (78) | 7 (70) |  |
| ART |  |  |  |  |  |  |  |  |  |  |
| TDF+3TC+DTG | 62 (75) | 23 (66) | 0.54 | 76 (70) | 9 (90) | >0.99 | 73 (74) | 7 (78) | 5 (50) | 0.31 |
| TDF+3TC+EFV | 19 (23) | 11 (31) |  | 29 (27) | 1 (10) |  | 24 (24) | 2 (22) | 4 (40) |  |
| Missing | 2 (2) | 1 (3) |  | 3 (3) | 0 (0) |  | 2 (2) | 0 (0) | 1 (10) |  |
| Placental malaria |  |  |  |  |  |  |  |  |  |  |
| None observed | 1 (1) | 1 (3) | 0.85 | 2 (2) | 0 (0) | 0.51 | 2 (2) | 0 (0) | 0 (0) | 0.43 |
| Active | 5 (6) | 1 (3) |  | 5 (5) | 1 (10) |  | 6 (6) | 0 (0) | 0 (0) |  |
| Active-chronic | 11 (13) | 6 (17) |  | 17 (16) | 0 (0) |  | 17 (17) | 0 (0) | 0 (0) |  |
| Past-chronic | 64 (77) | 26 (74) |  | 81 (75) | 9 (90) |  | 72 (73) | 8 (89) | 10 (100) |  |
| Missing | 2 (2) | 1 (3) |  | 3 (3) | 0 (0) |  | 2 (2) | 1 (11) | 0 (0) |  |

Abbreviations: T, term birth; PTB, preterm birth; LB, live birth (not neonatal death); PND, perinatal death; N, normal birth weight; LBW, low birth weight; M, missing birth weight; E, enrollment; D, delivery; ART, antiretroviral therapy; TDF, tenofovir; 3TC, lamivudine; DTG, dolutegravir; EFV, efavirenz
† All data are median (IQR) for continuous values or absolute numbers (percentages) for categorical values.
‡ Mann-Whitney test was used to analyze the relationship between the groups with numerical values, while Fischer's exact test was used for those without.
α Normal vs. low vs. missing birth weight. Kruskal-Wallis was used to analyze the relationship between all three groups with numerical values, a Fischer exact
test was used for those without, For comparison of birth weights, a Mann-Whitney test was performed.

### Plasma immune factor concentrations differ by anatomical location

To analyze concentration differences across placental, cord and postdelivery plasma, we grouped the 45 measured immune factors into four categories: chemokines, inflammatory cytokines, pro-/anti-inflammatory cytokines, or biomarkers (**Table S1**). Chemokine concentrations were highest in placental plasma: 13 of the 14 chemokines were significantly higher than in postdelivery plasma (**Figure 1a, S1)**, and 9 of the 14 were significantly higher than in cord blood plasma (**Figure 1b, S1)**. Conversely, postdelivery plasma exhibited the lowest overall chemokine concentrations (**Figure 1a,c,d,f** and **S1**). Inflammatory markers were also elevated in both postdelivery and placental plasma compared to cord blood (**Figure 1 b,c** and **S2**). Overall, we observed greater variation in immune factor concentrations across different anatomical compartments at delivery (cord, placental, and postdelivery plasma) than was observed across different maternal time points (**Figure 1**). The largest differences between placental plasma and both the cord blood plasma and postdelivery plasma were observed in the chemokines CCL2, CCL3, CCL4, and IL-8, as well as the inflammatory markers, IL-6, S100A8/S100A9, and SAA (**Figures S1-S3**, **Table S3**). Correlation matrices were generated to visualize associations between immune factors in cord and placental plasma (**Figures S5-S6**, **Tables S4-S5**). Few negative correlations were observed. In both placental and cord blood plasma, the strongest positive correlations were among MMP-9, MPO, and NGAL and S100A8/A9. Furthermore, placental plasma exhibited unique, strong positive correlations between IL-8, IL-1β, and IL-6.

### Immune factors associated with PTB

Next, we investigated whether immune factor levels were associated with PTBs. Several chemokines that have been shown to recruit T cells were associated with PTBs. CCL20 concentrations were higher at enrollment and postdelivery in PPLWH who gave birth prematurely, but was significantly lower in the placental plasma of PTBs compared to term births. Conversely, both CXCL9 and CXCL10 concentrations were significantly higher in the cord blood and postdelivery plasma of premature births (**Figure 2**, **Table S6**). Furthermore, gestational age at delivery was significantly negatively correlated with CCL20 and CXCL9 concentrations across various plasma types (**Figure S7**). MMP-2 was reduced in cord blood and postpartum plasma of PTBs versus term births. While MMP-9 and CST3 were also lower in PTB cord blood and placental plasma, both were significantly higher at enrollment (**Figure 2**, **Table S6**). All three markers significantly correlated with gestational age at delivery in at least one plasma type (**Figure S7**). IL-1RN, NGAL, and sTNFRSF1A concentrations were higher at enrollment in PPLWH who gave birth prematurely versus those who gave birth at term. A marker of skeletal and cardiac muscle damage, MB, was lower in the postdelivery and postpartum plasma of PTBs (**Figure S8**, **Table S6**). Because participants were enrolled at different times during their pregnancy, we performed a sensitivity analysis to determine if the timing of enrollment and corresponding plasma collection affected associations between PTB and immune factors. There were no significant differences between the levels of CST3 by trimester, despite this, PPLWH that enrolled in the second trimester and gave birth prematurely had significantly higher concentrations of CST3, this effect was not observed in PPLWH who enrolled in the third trimester (**Figure S9**).

**Figure 2.**
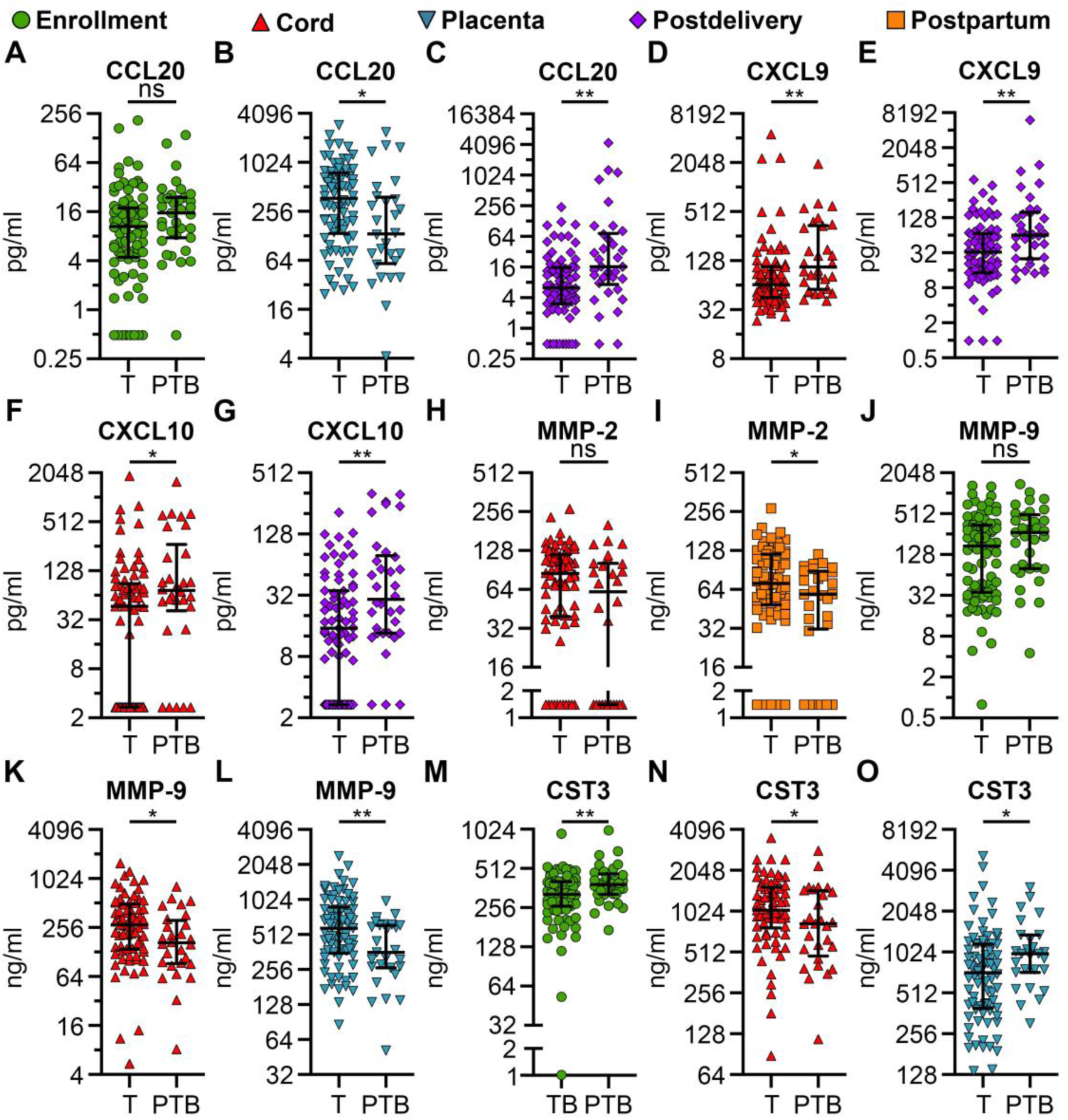
Immune factors associated with preterm birth. Immune factor concentrations were measured in plasma collected from participants at enrollment, postdelivery, and postpartum timepoints, as well as from placenta and cord blood. Concentrations were compared between individuals who delivered at term (T, n = 83) and from preterm births (PTB, n = 35). Each point represents an individual participant’s analyte concentration; horizontal lines indicate the median and IQR. Statistical significance was assessed using a two-tailed Mann–Whitney test. *p<0.05, **p<0.01, “ns” signifies 0.05<p<0.1. See **Table S6** for descriptive statistics.

### Immune factors associated with PND

Neutrophil-recruiting chemokines were generally higher in PND cases compared to live births. Specifically, CXCL1 was higher in both postdelivery and postpartum plasma, CXCL5 was higher only in postdelivery plasma, and IL-8 was higher in cord blood plasma. Markers of neutrophil and macrophage activation followed a similar pattern. MPO and NGAL concentrations were higher in the cord blood and placental plasma of PND cases, while S100A8/A9 was higher in both cord blood and postdelivery plasma. Furthermore, sCD14 was significantly higher only at enrollment, while significantly higher concentrations of sCD163 were observed in placental and postpartum plasma. A non-significant trend toward higher sCD14 persisted through postdelivery and postpartum periods. Dysregulation of L-VEGF and sVCAM-1, markers of endothelial activation and inflammation, were also evident; both markers were lower in cord blood plasma, whereas sVCAM-1 was lower in placental plasma and higher in postdelivery plasma in PND cases versus live births. (**Figure 3**, **Table S7**). IL-1RN and MB were higher in the placental and cord blood plasma of PND cases (**Figure S10**, **Table S7**). Trimester-specific analysis revealed that while CCL11 and CCL20 did not vary significantly across trimesters, both were significantly higher in PND cases enrolled during the third trimester. Conversely, CXCL9 and IGFBP4 varied significantly by trimester; CXCL9 remained significantly associated with PND in the third-trimester enrollment samples, and IGFBP4 showed a significant association with PND in the second trimester, but not the third. Conversely, sCD14 did not remain significant at enrollment when analyzed by trimester (**Figure S11**).

**Figure 3.**
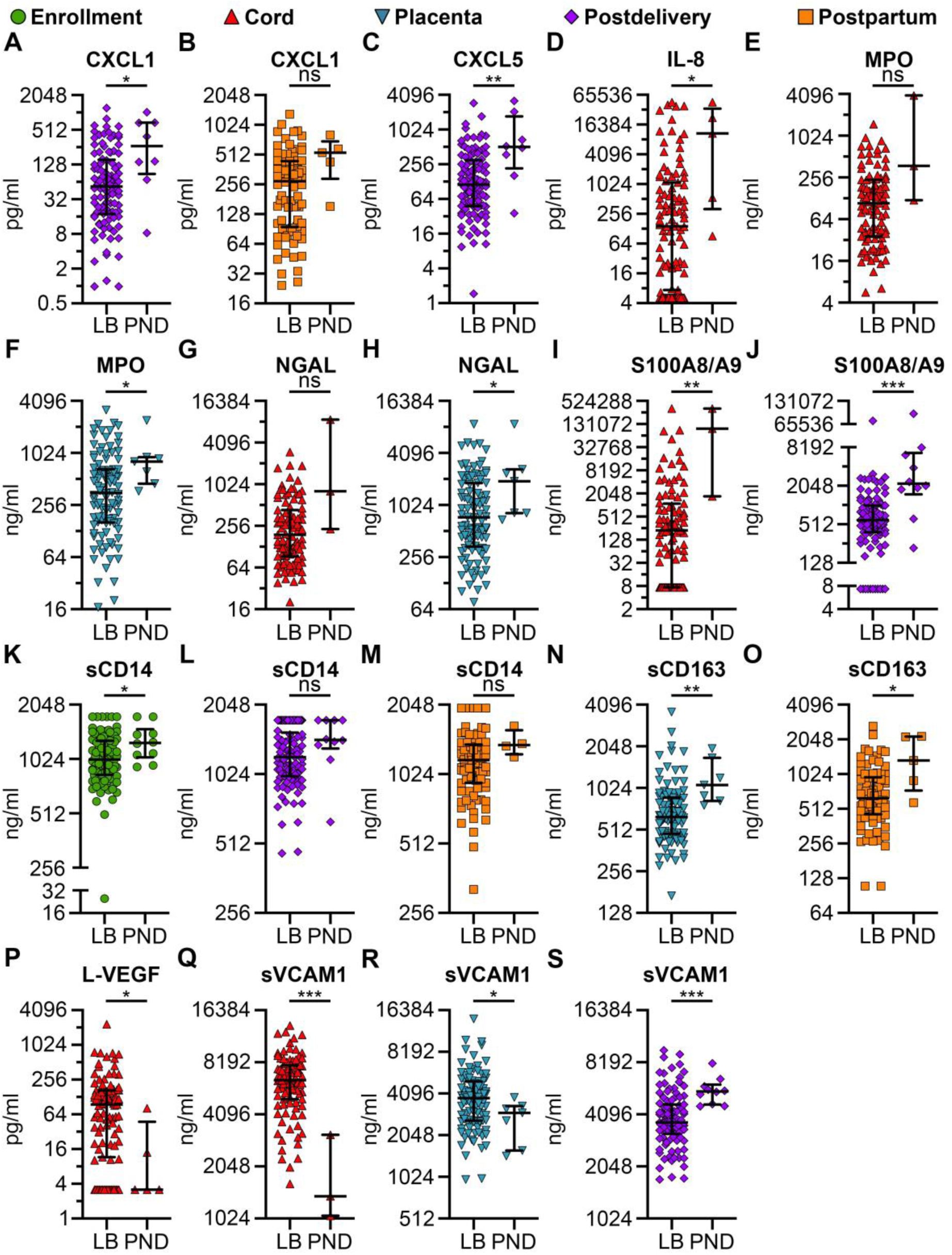
Immune factors associated with perinatal death. Immune factor concentrations were measured in plasma collected from participants at enrollment, postdelivery, and postpartum timepoints, as well as from placenta and cord blood. Concentrations were compared between live births (LB, n = 108) and perinatal deaths (PND, n = 10). Each point represents an individual participant’s analyte concentration; horizontal lines indicate the median and IQR. Statistical significance was assessed using a two-tailed Mann–Whitney test. *p<0.05, **p<0.01, ***p<0.001, “ns” signifies 0.05<p<0.1. See **Table S7** for descriptive statistics.

### Immune factors associated with LBW

LBW was associated with distinct inflammatory cytokine and chemokine profiles. CXCL9 and CXCL10 were higher in the postpartum plasma of LBW cases. At cenrollment, plasma concentrations of IL-18, S100A8/A9, and SPP1 were all higher in LBW cases, though SPP1 was not significantly so. Meanwhile, SAA concentration was more dynamic in LBW cases, being higher postdelivery and lower postpartum (**Figure 4** and **Table S8**). Furthermore, sICAM1 exhibited a more sustained association with LBW; concentrations were higher at enrollment and remained higher postdelivery (**Figure S12**), and it correlated with birth weight across various plasma types (**Figure S13**). Trimester-specific analysis revealed that both IL-18 and S100A8/A9 were significantly higher in PPLWH who enrolled in the third trimester and subsequently gave birth to a LBW neonate (**Figure S14**). In a separate analysis of individuals lacking birth weight data, only six immune factors showed significant results (**Figure S15**, **Table S8**).

**Figure 4.**
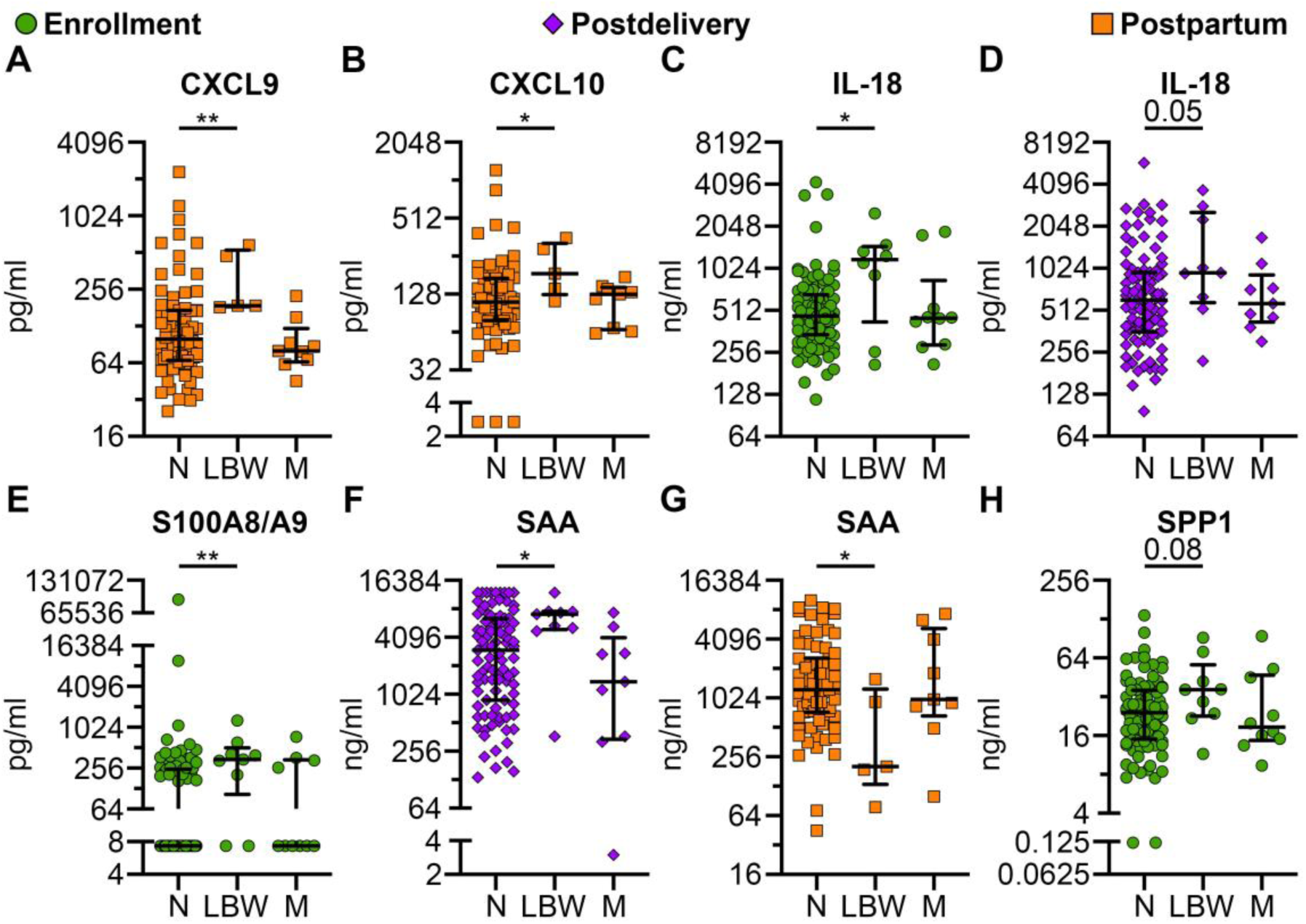
Immune factors associated with low birthweight. Immune factor concentrations were measured in plasma collected from participants at enrollment, postdelivery, and postpartum timepoints. Concentrations were compared between individuals with normal birthweight (N, n = 99), low birthweight (LBW, n = 9), and those with missing birthweight data (M, n = 10). Each point represents an individual participant’s analyte concentration; horizontal lines indicate the median and IQR. Statistical significance tested by Kruskal-Wallis uncorrected for multiple comparison. *p<0.05, **p<0.01, p-values between 0.05 and 0.1 labeled. No statistically significant difference was found between N and M. See **Table S8** for descriptive statistics.

### Composite inflammatory score across birth outcomes

The composite inflammatory score was higher in all adverse outcomes compared to the no adverse outcome group. The highest inflammatory score was associated with PND, with the inflammatory score primarily driven by immune factors present in the umbilical cord and postdelivery plasmas. Postpartum plasma was primarily a negative driver of the inflammatory score (i.e. was less inflammatory) in the no adverse outcome, PTB, and PND groups (**Figure 5**). Notably, this negative contribution from postpartum plasma was absent in LBW cases (**Figure 5)**, where multiple inflammatory cytokines were observed postpartum (**Figure 4)**.

**Figure 5.**
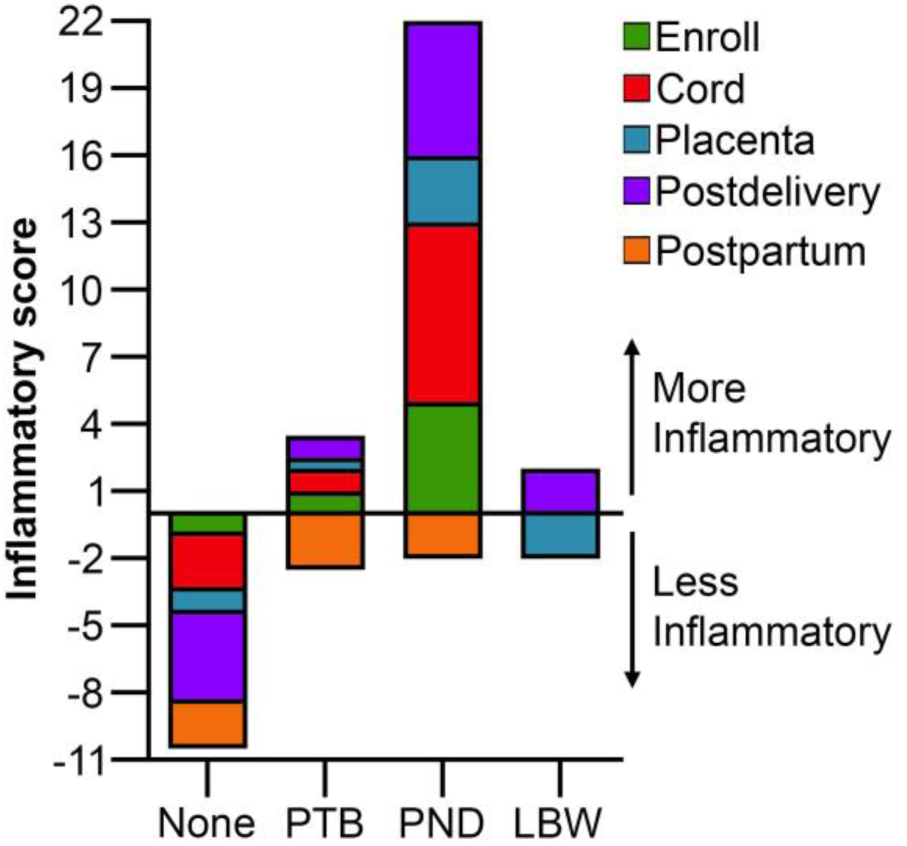
Composite inflammatory score across birth outcomes. Stacked bars represent the cumulative median score for each group, summed across five plasma and tissue sources. Groups include: no known adverse outcome (None), preterm birth (PTB), perinatal death (PND), and low birthweight (LBW). Note that participants can belong to multiple groups, and those individuals who were missing birthweight data but had no other adverse outcome were excluded from this analysis. Positive values indicate a more inflammatory profile. Scores were derived from 26 pro-inflammatory and 3 anti-inflammatory factors (**see Table S9**) using quartile-based calculations (**Table S3**).

## Discussion

Our study of pregnant people living with HIV revealed several key findings regarding the maternal-fetal immune environment at delivery. Overall immune factors were enriched in placental plasma compared to cord blood or postdelivery plasma, despite all samples being collected at or near delivery. When examining adverse outcomes, dysregulated chemokines and biomarkers were most frequently associated with PTB. Although fewer immune factors were associated with LBW overall, multiple markers were significantly associated with this outcome at enrollment, highlighting their potential as early predictive biomarkers. In contrast, PND was associated with widespread immune alterations, particularly within cord blood, placental, and postdelivery plasma. Finally, while the composite inflammatory score was higher for all adverse outcomes relative to individuals with no known adverse outcome, the overall level of inflammation remained lower than expected—with the notable exception of PND cases. These cases exhibited the greatest inflammatory burden driven primarily by postdelivery and cord blood plasma.

Taken together, these findings highlight that both anatomical and temporal distribution of immune factors matters just as much as their overall concentrations.

To understand why placental plasma showed such a distinct and highly concentrated immune factor profile, we first considered its unique biological context. Placental and cord blood plasma are rarely utilized in pregnancy research compared to maternal plasma [13,21,78–81], likely due to collection challenges and limited collection windows in humans. Placental blood collection can be achieved through various methods, yielding different admixtures of fetal and maternal blood. The method used in this study has previously been shown to produce primarily maternal blood from the intervillous space [65].

However, as the maternal blood passes by the placenta it is enriched for factors secreted by the placenta itself. Previous studies have demonstrated that isolated placental trophoblasts and placental explants secrete immune factors *in vitro* and *ex vivo* [82–86]. Additionally, the placenta produces hormones, including immunomodulatory hormones such as cortisol [87–90]. This complexity makes it difficult to distinguish immune factors that are maternally derived from those secreted by the placenta or induced in response to placental hormones, particularly across the dynamic transition from parturition to postpartum [88–90]. Furthermore, few studies have examined changes in immune profiles from onset of labor through the early postpartum period [91–94], further complicating interpretation. Our longitudinal data shows that most chemokines, including CCL2 and CCL17 decreased from enrollment to postdelivery [76], yet remained higher in the placental plasma at delivery. In contrast, chemokines such as CCL3 remain low in maternal circulation at both enrollment and postdelivery time points [76], but are higher in placental plasma, indicating at least transient placental expression at delivery. Taken altogether, these *in vitro, ex vivo* and *in vivo* findings suggest that the placenta may secrete certain immune factors, including chemokines, and that these may end up in maternal circulation during pregnancy. Given the central role in fetal development and the essential functions it performs [95], the placenta has long been recognized as an important determinant in adverse fetal outcomes [16,24,25,96]. In rare cases, an abnormal placenta can even lead to maternal death [96]. Our data raise the possibility that placental dysregulation may also have subtler effects on maternal physiology through immune signaling into the maternal circulation.

In our cohort, the primary drivers of this placental dysregulation were likely infectious. Both HIV and *Plasmodium* infections can cause placental dysfunction and chronic placental inflammation [16–18,97]. Because all participants were living with HIV, we were unable to directly assess the contribution of HIV itself; however, analysis of immune factors in placental plasma showed no significant differences between those with a detectable and undetectable viral load. Similarly assessing whether placental malaria contributed to immune factor levels was limited by the small number of placentas without evidence of infection. Nevertheless, three immune factors significantly correlated with placental malaria: IFN-λ-1, sCD14, and MB (**Figure S16)**. Previous data has demonstrated constitutive expression of IFN-λ-1 from placental epithelial cells [82,85], making its unexpectedly low levels in our study notable. A previous study demonstrated a decrease in IFN-λ-2 in response to chronic malarial infection during pregnancy [98] suggesting placental malarial infection may similarly suppress IFN-λ-1 expression. Since, IFN-λ-1 is generally considered immunomodulatory [99], its reduced expression may permit an overall increase in inflammation.

The near-universal presence of placental malaria that limited these immune comparisons is perhaps less surprising given the regional context and time frame of our study. The WHO reports that the DRC accounted for 12% of the world’s reported malaria cases in 2021, second only to Nigeria [100]. Previous studies in African populations have reported high subregional prevalence alongside lower pooled regional rates [101]. Our cohort enrolled from October 2020 to May 2021; notably, the DRC experienced a decrease in testing during this period due to the ongoing COVID-19 pandemic [102,103], which may have resulted in underdiagnosis and undertreatment, potentially allowing ongoing transmission to go undetected in the broader community. While earlier research indicates that individuals in their first or second pregnancies are most susceptible to placental malaria [41–43], a number of studies in pregnancies affected by HIV without ART suggest that gravidity is less of a factor in placental invasion in the context of HIV-*Plasmodium* coinfection [101,104,105]. Although PPLWH in our study were on ART and we had a large number of multigravida individuals, our placental pathologists diagnosed all but two available placentas as having evidence of current or past *Plasmodium* invasion of the placenta. Correspondingly, we observed a large number of stillbirths in our study (7% versus a national average of 2.6% [106]) and PTBs (30% versus a national average of 12.4% [107]). While HIV can increase the risk of both stillbirth and PTB [27,28,30,32,37,38], *Plasmodium* elevates these risks as well [37,39,108–110], particularly if parasites infect the placenta [39,97,109,110]. Consequently, this exceptionally large burden of placental malaria could help to explain the observed increases in stillbirths and PTB within our cohort.

While malaria likely contributed to specific adverse outcomes, perinatal death more broadly remains a critical yet understudied phenomenon [111–114], partly due to inconsistent terminology [115–117] and psychosocial stigma that may lead to underreporting [112,118,113]. Known risk factors include: infection, placental and umbilical cord pathology, birth weight, gestational age at delivery, and maternal history/health [24,31,32,115,119–121]. PND cases in our cohort primarily exhibited higher concentrations of chemokines linked to T cell, monocyte and granulocyte recruitment [122], as well as, higher levels of markers of neutrophil and macrophage activation [123–127]. Both neutrophils and macrophages have been associated with PND in placental, cord, and/or fetal tissues, though neutrophil activation in particular is usually associated with infection especially in the fetal compartment [16,25,26,128]. These findings suggest that innate immune cells may play an important role in the immunopathology of PND. Beyond the pathways specific to PND, we also identified a distinct subset of immune factors including: CXCL9, IL-1RN, MB, NGAL, S100A8/A9 and sCD163 that were not associated with a particular adverse outcome, but rather served as shared markers of multiple adverse outcomes. Although these are not necessarily a part of a particular pathway, they serve common roles as biomarkers of inflammation and tissue damage. For example, NGAL, S100A8A9 and sCD163, which are secreted by and cleaved from activated neutrophils and macrophages [123,126,129], were more highly expressed in cord blood and/or placental plasma of PND cases. Both S100A8/A9 and NGAL can sequester and regulate metals to affect antimicrobial activity [123,130], but S100A8/A9 also acts as a damage associated molecular pattern that can increas inflammation [131].

Among these shared markers of tissue damage, we identified myoglobin, a marker of striated muscle injury [132,133], as being associated with multiple adverse outcomes. Myoglobin was higher in PND cases, indicating possible skeletal or cardiac injury in the fetal compartments. This difference was observed both in stillbirths and neonatal deaths, suggesting the damage is severe, but not uniformly fatal prior to delivery. In contrast, PTB was associated with decreased myoglobin levels postdelivery despite no difference in delivery mode; this decrease potentially reflects a shorter second-stage labor—the period of maximum muscle strain—which has previously been shown to be briefer in PTBs compared to term births [134,135]. Alternatively, given the rapid clearance of myoglobin from circulation after acute injury [136] differences in sampling timing could account for this finding, unfortunately in this study the timing of the postdelivery blood draw was not recorded.

While myoglobin highlights physical tissue damage, specific chemokines pointed to shared inflammatory signaling across multiple adverse outcomes. Most notable was CXCL9, a chemotactic factor primarily responsible for recruiting T cells to inflammatory sites [122]. In pregnancy, CXCL9 has been associated villitis of unknown etiology, an inflammatory placental disease linked to PTBs, LBW, and PND [16,26,137,138]. While a previous study suggested that CXCL9 may be predictive of both pregnancy loss and preterm delivery [139], our analysis found CXCL9 is predictive of PND and associated with PTB. Interestingly, CXCL11, a chemokine that signals through the same receptor as CXCL9, was lower in the placental plasma of PND cases, an opposite trend to CXCL9. A similar phenomenon has been observed in the placentas of spontaneous versus elective abortions [140]. *In vitro* studies have demonstrated that CXCL11 is a stronger agonist for CXCR3 than CXCL9 and is better at internalizing the receptor. Additionally, CXCL11 has been shown to preferentially drive T cell polarization toward a more regulatory phenotype, whereas CXCL9 drives polarization toward a more inflammatory Th1 or Th17 subset [141]. CXCL9, but not the related CXCL10 and CXCL11, is a broad marker of adverse pregnancy outcomes across multiple maternal time points and anatomic locations.

In stark contrast to the pro-inflammatory profile of CXCL9, we also observed concurrent elevations of IL-1RN. This potentially anti-inflammatory molecule competitively inhibits the pro-inflammatory cytokines IL-1α and IL-1β [142]. Because of this, multiple drugs have been developed to block IL-1 signaling [143–145]. One preclinical drug, Rytvela significantly reduced not only PTB, but also fetal death in a mouse model of LPS-mediated PTB [146]. In our study, IL-1RN was consistently higher in multiple plasma types across adverse outcomes was IL-1RN; however, we did not observe significant differences in IL-1β under any conditions. Given that IL-1β and IL-1RN release occur through different mechanisms this is not surprising [147]. We did not measure IL-1α, so it is possible that it is higher in adverse outcomes, however, it is equally plausible that IL-1RN release was induced independent of other IL-1 family members.

## Limitations

While this pilot study provides valuable preliminary insights, several limitations should be noted. Methodologically, our study was conducted within a larger cohort of pregnant people living with HIV and lacked an HIV-negative pregnant comparator group. Additionally, as previously discussed [76], we did not control for the timing of enrollment or record the specific timing of the postdelivery time point. Finally, because this is a pilot study, our sample size is limited and drawn from a concentrated geographic area. Given that immunologic signatures in people living with HIV and pregnant individuals are known to vary regionally [148,149], the specific signatures observed here may not be broadly generalizable. Ultimately, these constraints highlight a clear path forward, emphasizing the need for larger, geographically diverse cohorts to validate and expand upon these initial findings.

## Conclusions

Ultimately, this study demonstrated that pregnant people living with HIV on ART exhibit distinct, compartment-specific soluble immune profiles that are significantly associated with adverse outcomes. By analyzing plasma in multiple anatomical compartments across the peripartum, we established that inflammation is not uniform across the fetal-maternal interface. We further demonstrated that there were both anatomical and temporal-specific immune factors significantly associated with adverse outcomes, which suggests that despite ART, adverse outcomes in PPLWH and placental malaria are driven by complex, localized immune system dysregulation. This highlights the critical need to test whether anti-inflammatory therapies could improve pregnancy oucomes among pregnant people living with HIV taking antiretroviral therapy, particularly those that are also impacted by placental malaria.

## Abbreviations

3TC: Lamivudine
APP: Acute phase protein
ART: Antiretroviral therapy
C: Cord blood
CQI: Continuous Quality Improvement
CQI-PMCT: Continuous Quality Improvement-Prevention of Mother to Child Transmission
DAMP: Damage-associated molecular pattern
DTG: Dolutegravir
E: Enrollment
EFV: Efavirenz
HIV: Human immunodeficiency virus
IQR: Interquartile range
LB: Live birth
LBW: Low birthweight
LoD: Limit of detection
M: Missing birthweight
N: Normal birth weight
P: Placental blood
PD: Postdelivery
PND: Perinatal death
PP: Postpartum
PPLWH: Pregnant people living with HIV
PTB: Preterm birth
T: Term birth

## Declarations

### Ethics approval and consent to participate

The authors confirm that the ethical policies of the journal, as noted on the journal’s author guidelines page, have been adhered to and the appropriate ethical review committee approval has been received. The study conformed to the US and DRC Policies for the Protection of Human Subjects. This is an observational cohort study within a larger study approved by Institutional Review Boards (IRB) at The Ohio State University (IRB study ID: 2015H0440), Albert Einstein College of Medicine (protocol #2020-12018) and the Kinshasa School of Public Health Ethics Committee (protocol #0001 1–04101-00001365292–20). Written informed consent was obtained directly from participants. Data were de-identified prior to processing and analysis.

### Consent for publication

Not applicable

### Availability of data and materials

All data needed to evaluate this manuscript are in supplemental tables.

### Author contributions

Conception and overall study design: JJK, MY, PB, MT, JC. Acquisition of clinical samples: MT, FLK, BLM, PB. Placental pathology interpretation: FLK, BLM. Acquisition/interpretation cytokine data: JC. Drafting manuscript: JC. Revising for intellectual content: JC, NZ, NTF, MY, JJK, JAM. Final approval of completed manuscript: JC, NZ, MT, FLK, BLM, PB, JAM, NPML, NTF, MY, JJK.

## Acknowledgments

We thank all of the participants who contributed to the findings of this study. We acknowledge the contribution of the following site investigators of the CQI-PMTCT study team: Godelive Aitikalema, Ali Alisho, Elysée Bayayana, Fabrice Bumwana, Pierre Dianzenza, Jean Claude Dinanga, Georges Kihuma, Willy Lukumu, Fidèle Lumande, Zouzou Masevo, Fanny Matadi, Rachel Mushiya, Marie Therèse Mwela, José Nlandu, Pearl Tenatena, Marie Tshibuabua, Bienvenu Kawende and Noro LR Ravelomanana. We are grateful to participating clinics, provincial and national health authorities. We also acknowledge the support we have received from the administrative staff of The Ohio State University, Albert Einstein College of Medicine and Kinshasa School of Public Health. In addition, we acknowledge the efforts and support of Christina Cotrone, Brian Reinert and Kate Ailstock.

This research was supported by the President’s Emergency Plan for AIDS Relief (PEPFAR), and the National Institutes of Health (R01HD087993, R01HD05526, U01AI096299). The funders had no role in study design, data collection, data analysis and interpretation, preparation of the manuscript, or decision to submit.

---

**Figure S1.**
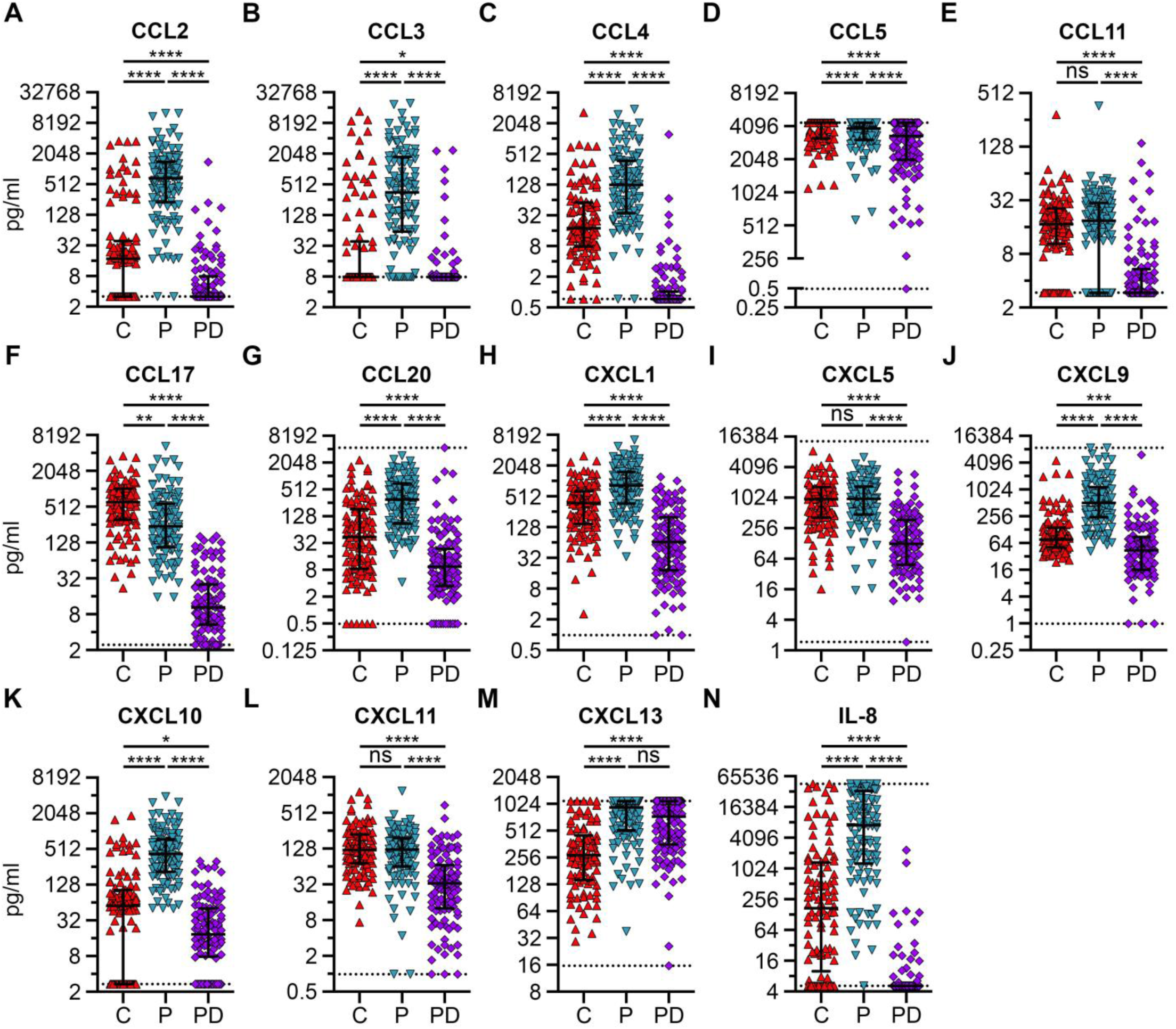
Chemokine expression in cord, placental, and postdelivery plasma. Chemokines were measured using (A-L, N) LEGENDplex assays and (M) ELISAs in plasma collected from cord (C) blood, placental (P) blood, and at the postdelivery (PD) timepoint. Each point represents an individual participant’s analyte measurement, with horizontal lines indicating the median and IQR. Dotted lines represent limits of detection (LoD), see **Table S2** for all LoD values and **Table S3** for descriptive statistics. Statistical significance tested by Kruskal-Wallis with a Dunn’s correction for multiple comparison. ns=not significant, *p<0.05, **p<0.01, ***p<0.001, ****p<0.0001.

**Figure S2.**
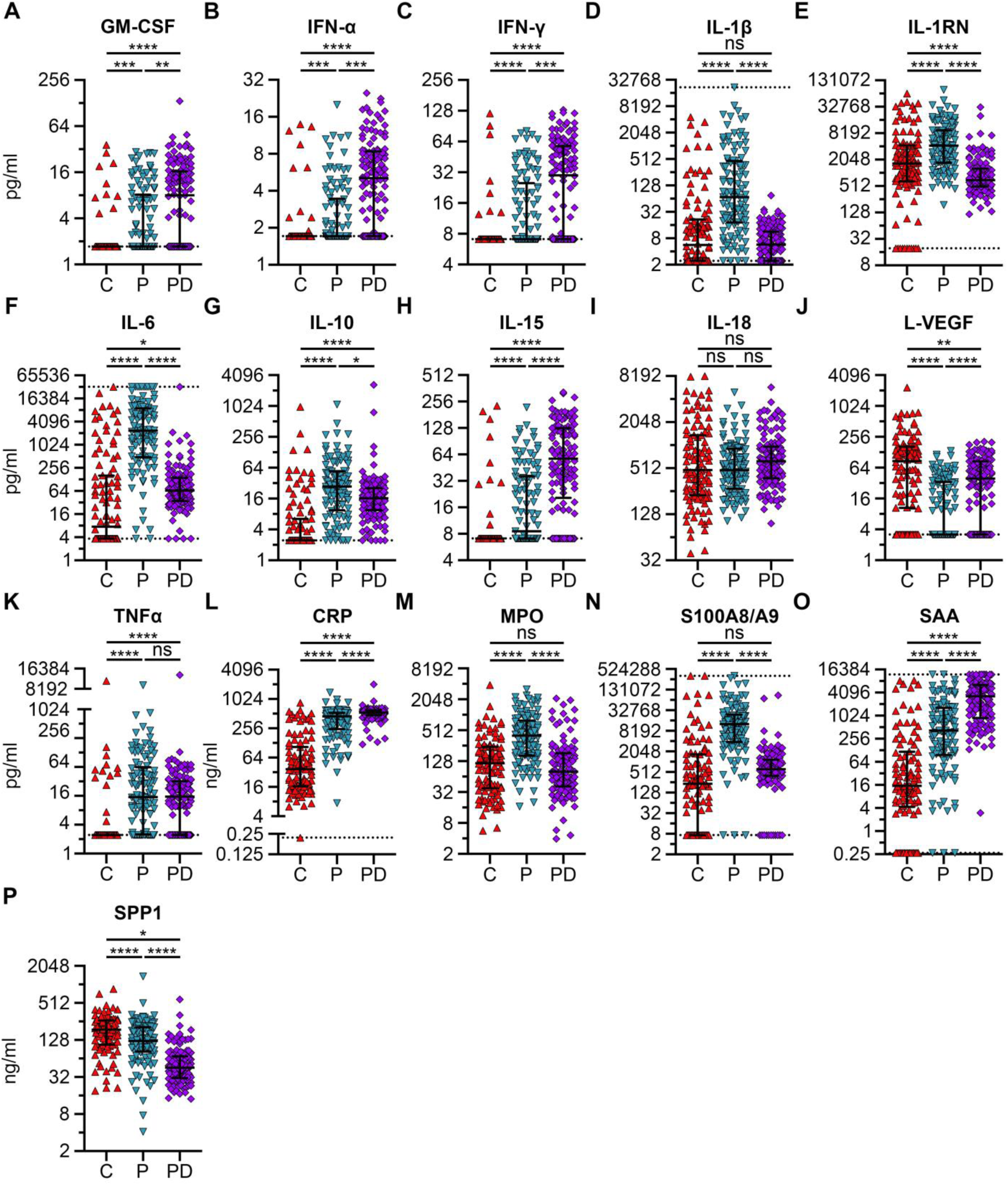
Pro/anti-inflammatory expression in cord blood, placental, and postdelivery plasma. Pro- and anti-inflammatory cytokines were measured using LEGENDplex assays in plasma collected from cord blood (C), placental (P) blood, and at the postdelivery (PD) timepoint. Immune factors were graphed in pg/ml (A-K) or ng/ml (L-P). Each point represents an individual participant’s analyte measurement, with horizontal lines indicating the median and IQR. Dotted lines represent LoD, see **Table S2** for all LoD values and **Table S3** for descriptive statistics. Statistical significance tested by Kruskal-Wallis with a Dunn’s correction for multiple comparison. ns=not significant, *p<0.05, **p<0.01, ***p<0.001, ****p<0.0001.

**Figure S3.**
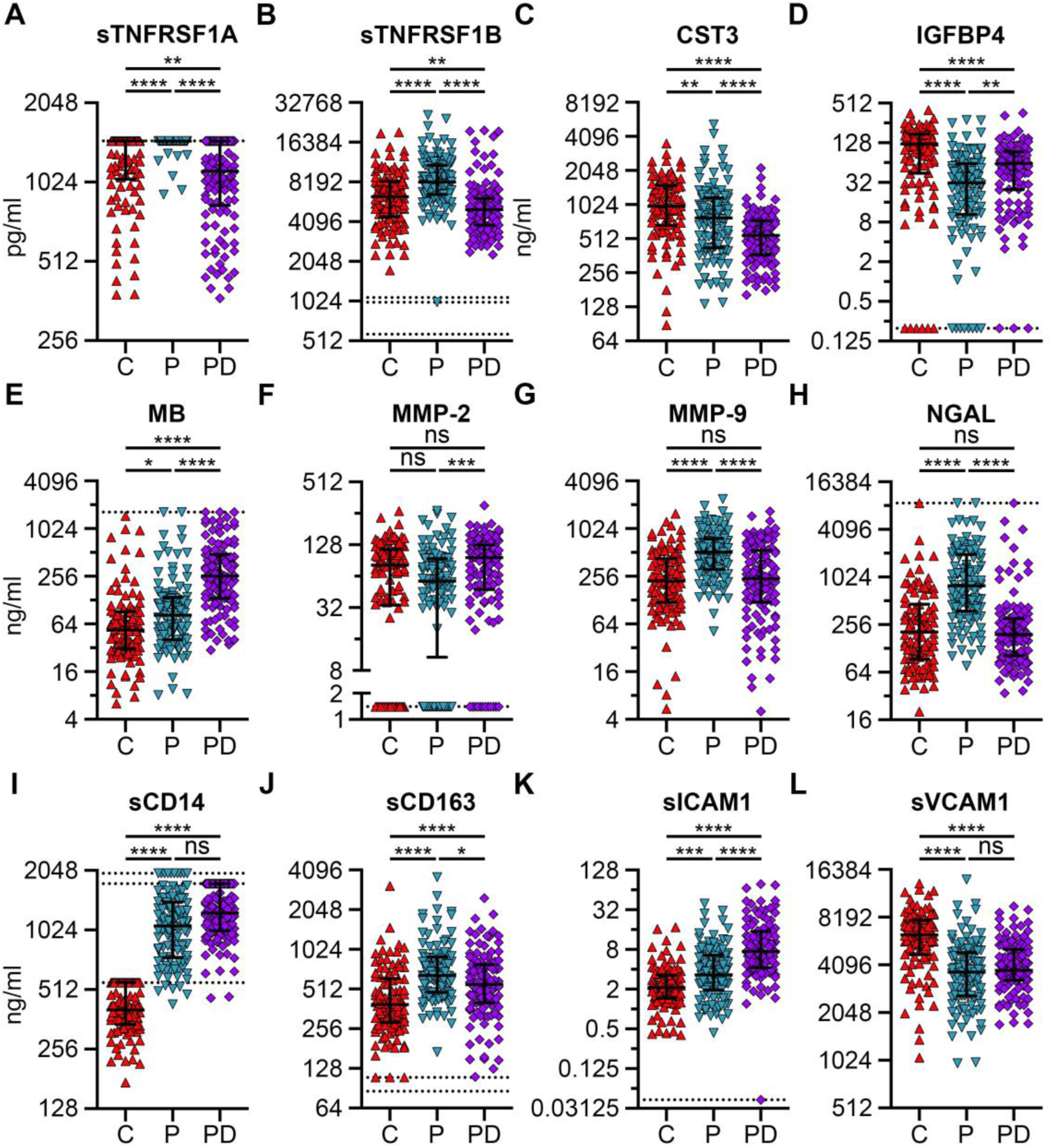
Soluble factor expression in cord, placental, and postdelivery plasma. Soluble factors were measured using (A-B, I-J) ELISAs and (C-H, K-L) LEGENDplex assays in plasma collected from cord (C) blood, placental (P) blood, and at the postdelivery (PD) timepoint. Immune factors were graphed in pg/ml (A-B) or ng/ml (C-L). Each point represents an individual participant’s analyte measurement, with horizontal lines indicating the median and IQR. Dotted lines represent LoD, see **Table S2** for all LoD values and **Table S3** for descriptive statistics. Statistical significance tested by Kruskal-Wallis with a Dunn’s correction for multiple comparison. ns=not significant, *p<0.05, **p<0.01, ***p<0.001, ****p<0.0001.

**Figure S4.**
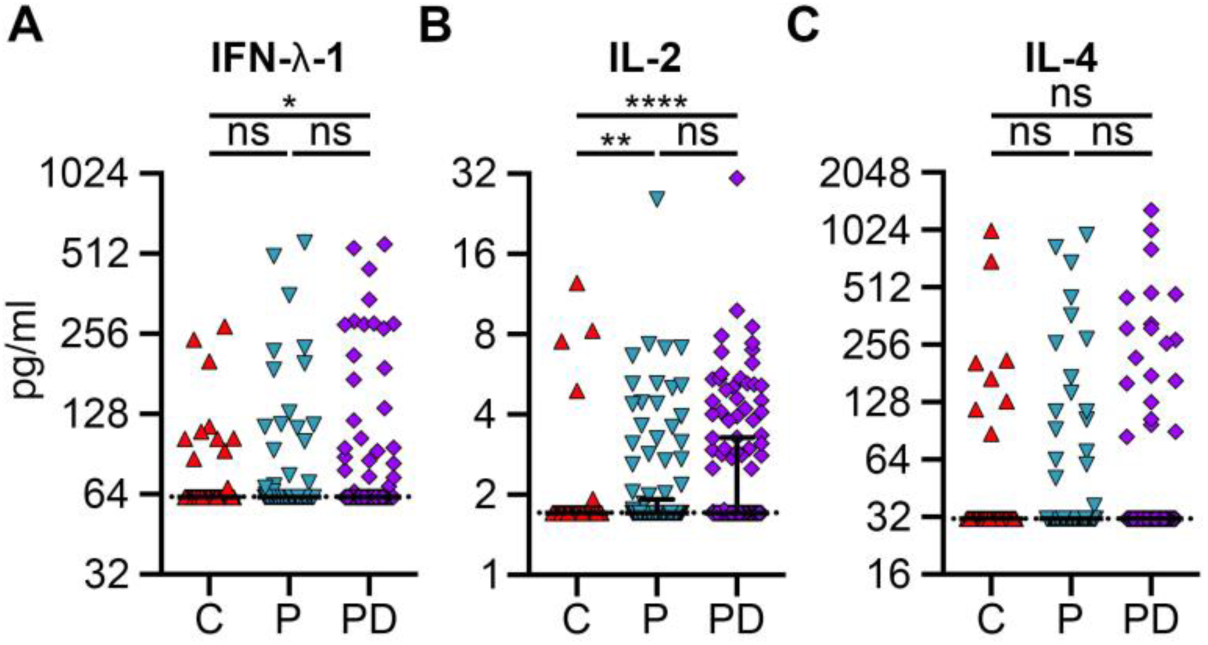
Immune factors with a high proportion of values below the LoD across all plasma types. Immune factors were measured using (A, C) ELISAs and (B) LEGENDplex assays in plasma collected from cord (C) blood, placental (P) blood, and at the postdelivery (PD) timepoint. Each point represents an individual participant’s analyte measurement, with horizontal lines indicating the median and IQR. Dotted lines represent LoD, see **Table S2** for all LoD values and **Table S3** for descriptive statistics. Statistical significance tested by Kruskal-Wallis with a Dunn’s correction for multiple comparison. ns=not significant, *p<0.05, **p<0.01, ****p<0.0001.

**Figure S5.**
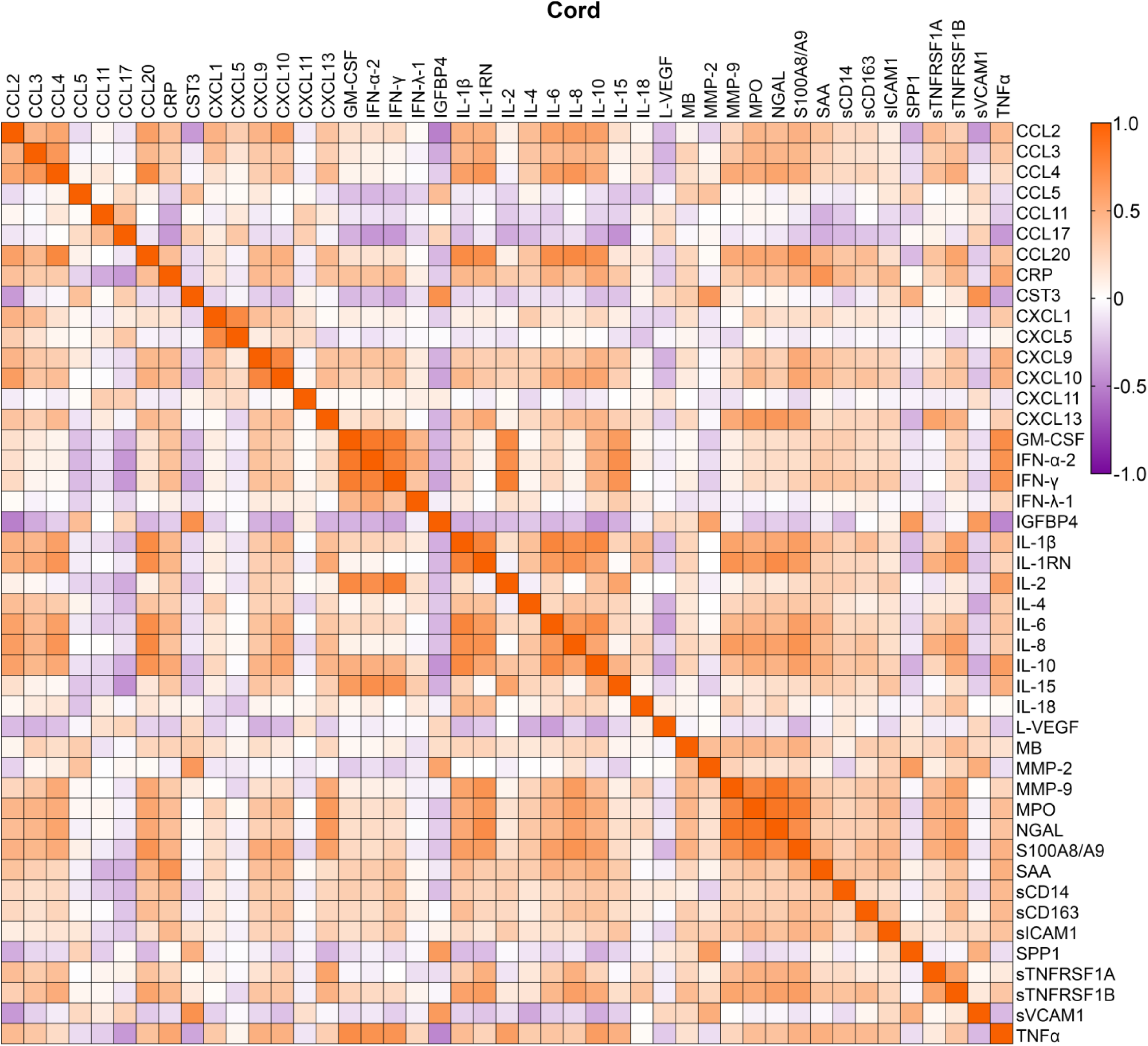
Correlations between immune factors in cord blood plasma. Graphical representation of two-tailed Spearman correlations between immune factors in fetal cord blood plasma. Spearman r and p-values are in **Table S4**.

**Figure S6.**
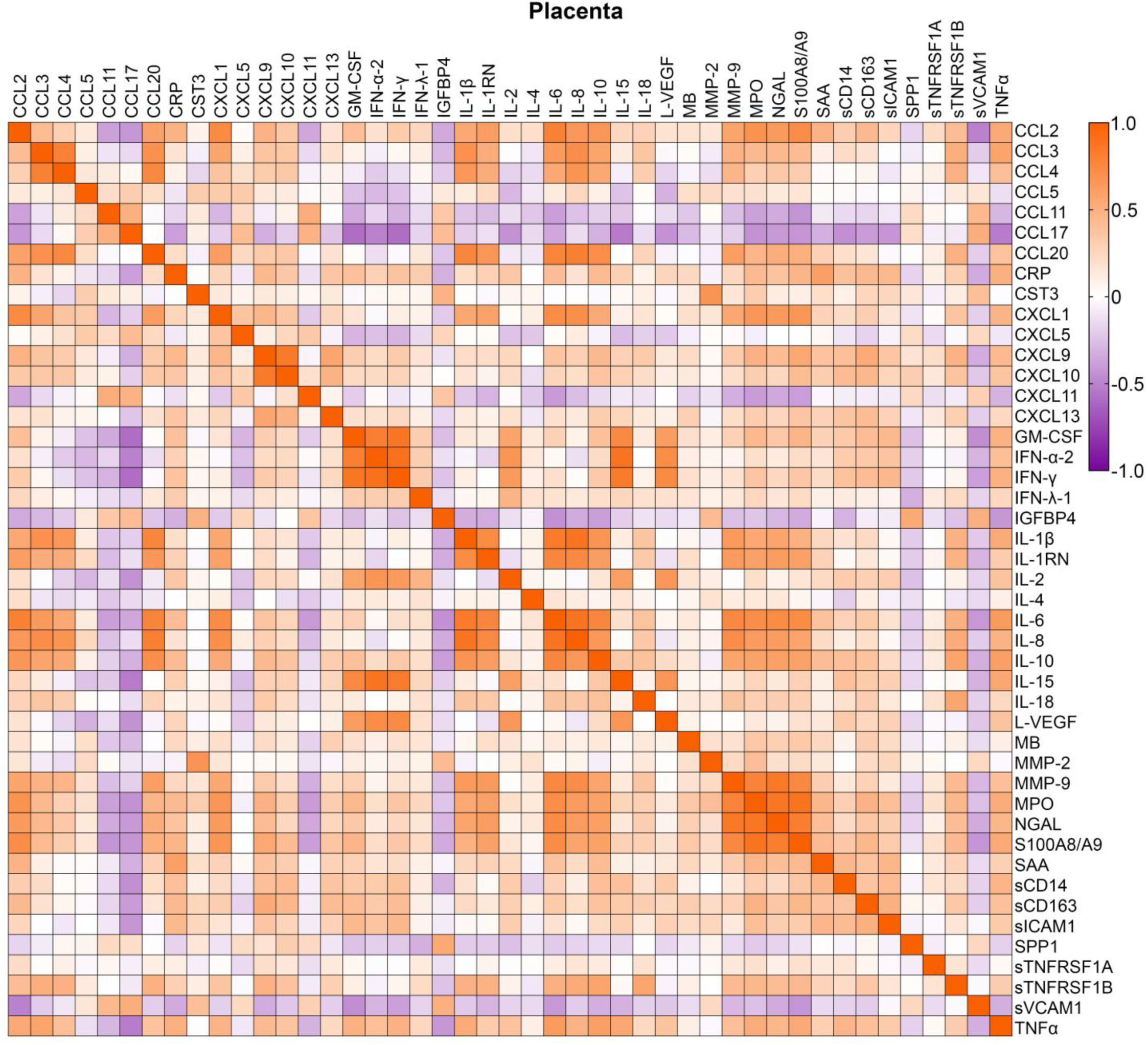
Correlations between immune factors in placental plasma. Graphical representation of two-tailed Spearman correlations between immune factors in placental plasma. Spearman r and p-values are in **Table S5**.

**Figure S7.**
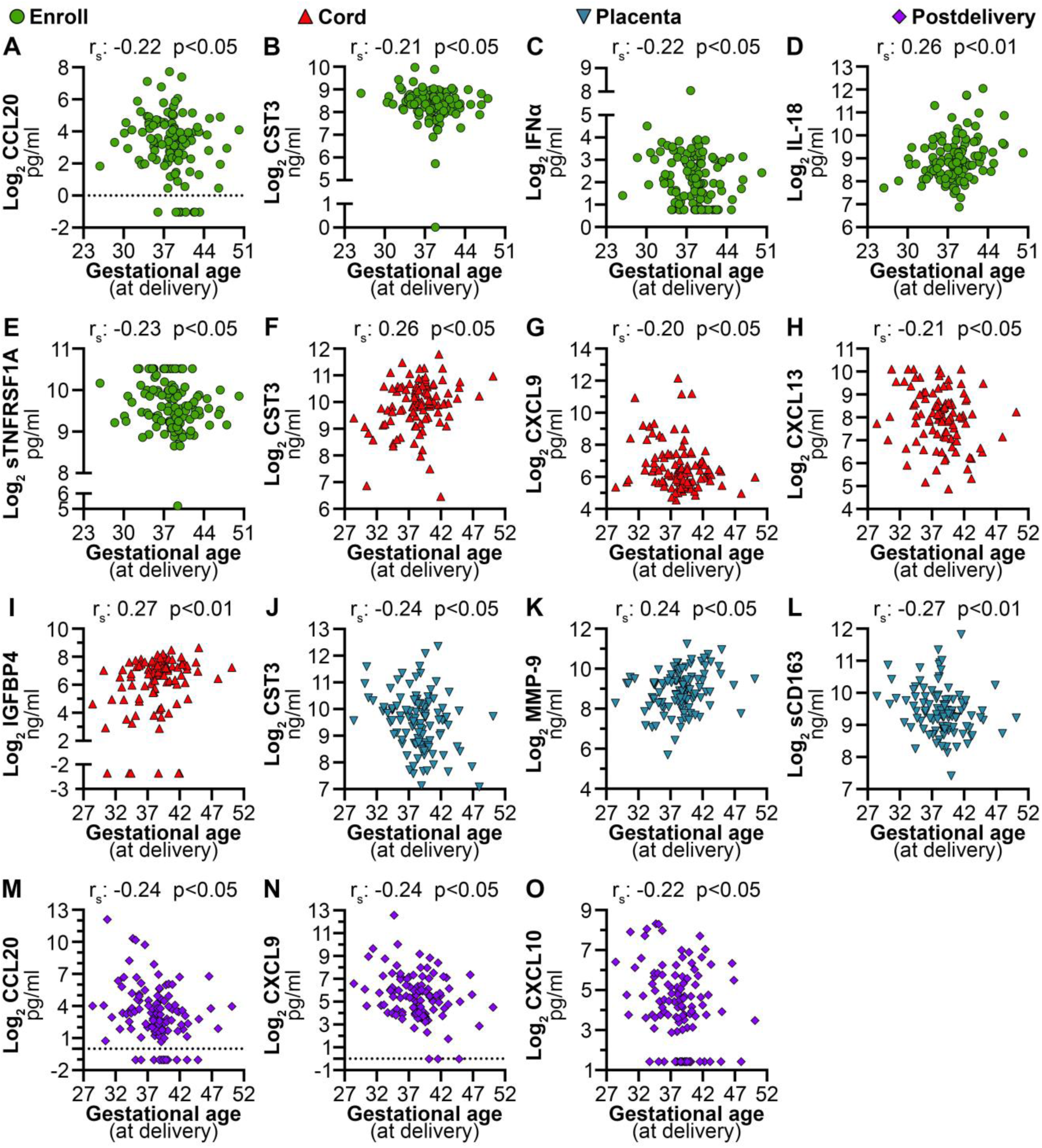
Immune factors that significantly correlate with gestational age at delivery. Log_2_ concentration of immune factors were plotted against the gestational age at delivery in weeks. Each dot represents an individual participant’s analyte measurement. Statistical significance tested by Spearman correlation (r_S_) and approximate p-value of the correlation displayed. Only those with a p-value<0.05 and an r_S_>0.2 or <-0.2 were considered significant.

**Figure S7.**
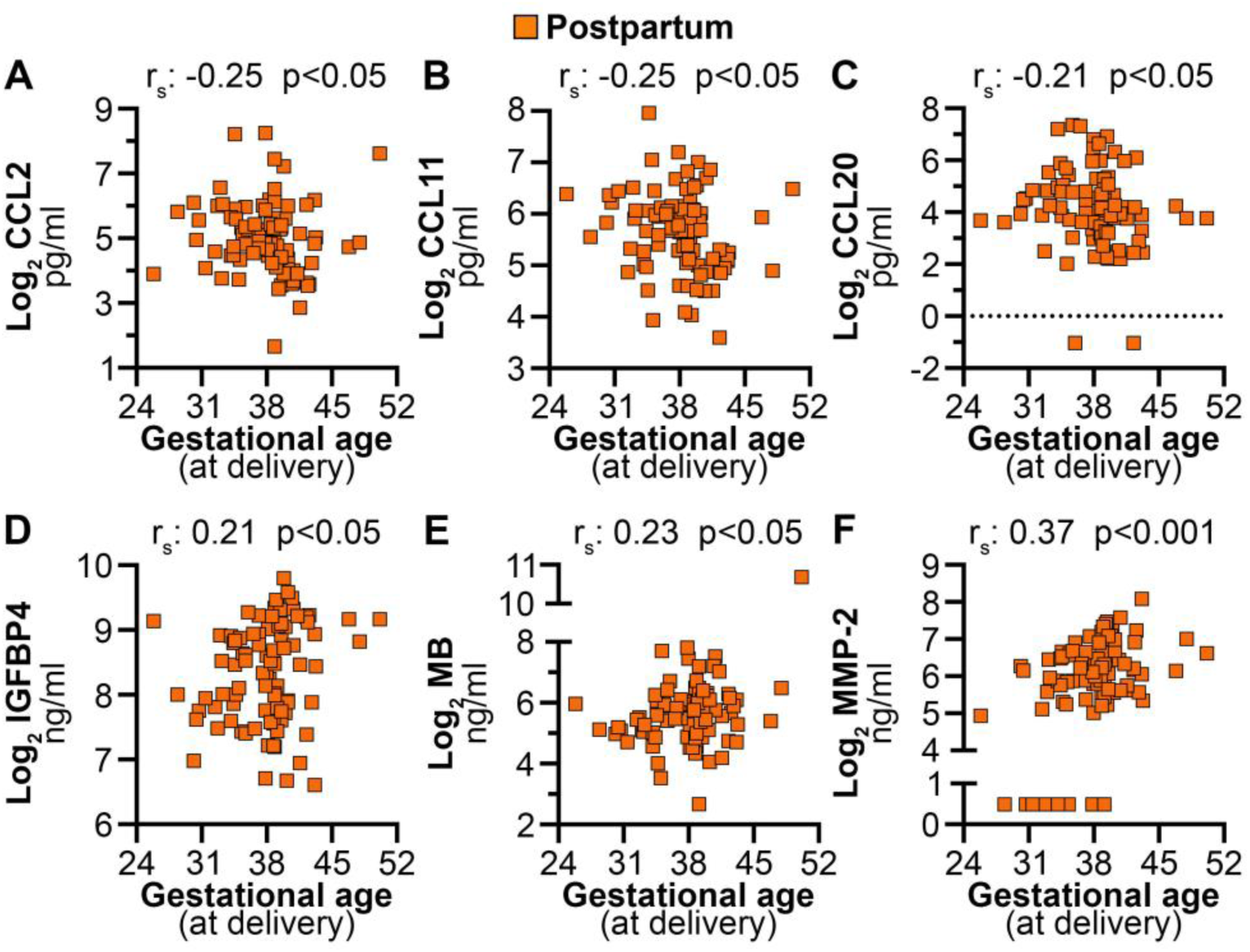
continued. Immune factors that significantly correlate with gestational age at delivery. Log_2_ concentration of immune factors were plotted against the gestational age at delivery in weeks. Each dot represents an individual participant’s analyte measurement. Statistical significance tested by Spearman correlation (r_S_) and approximate p-value of the correlation displayed. Only those with a p-value<0.05 and an r_S_>0.2 or <-0.2 were considered significant.

**Figure S8.**
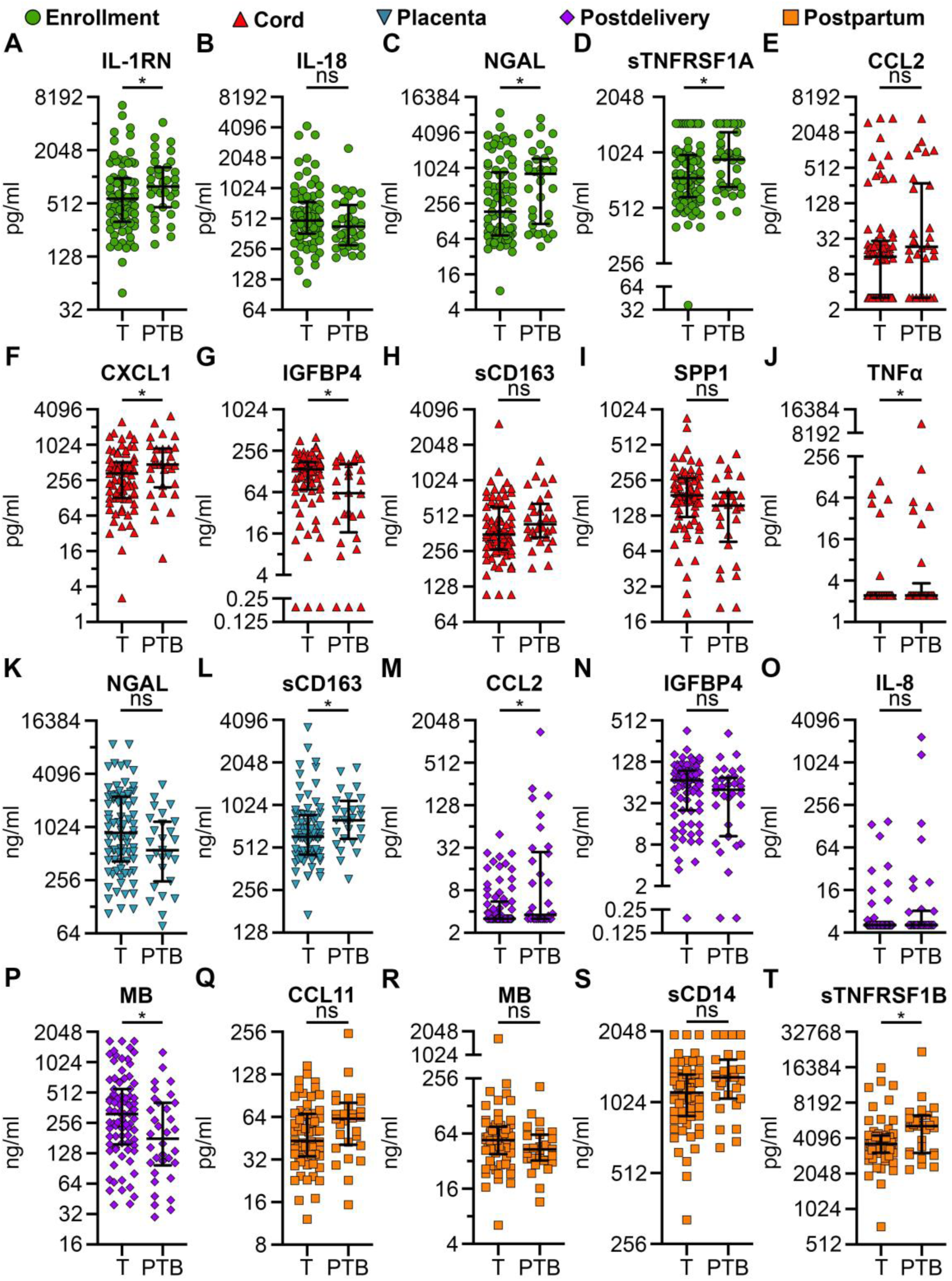
Immune factors associated with preterm birth. Immune factor concentrations were measured in plasma collected from participants at enrollment, postdelivery, and postpartum timepoints, as well as from placenta and cord blood. Concentrations were compared between individuals who delivered at term (T, n = 83) and preterm births (PTB, n = 35). Each point represents an individual participant’s analyte concentration; horizontal lines indicate the median and IQR. Statistical significance was assessed using a two-tailed Mann–Whitney test. *p<0.05, ns signifies a p value between 0.05 and 0.1. See **Table S6** for descriptive statistics.

**Figure S9.**
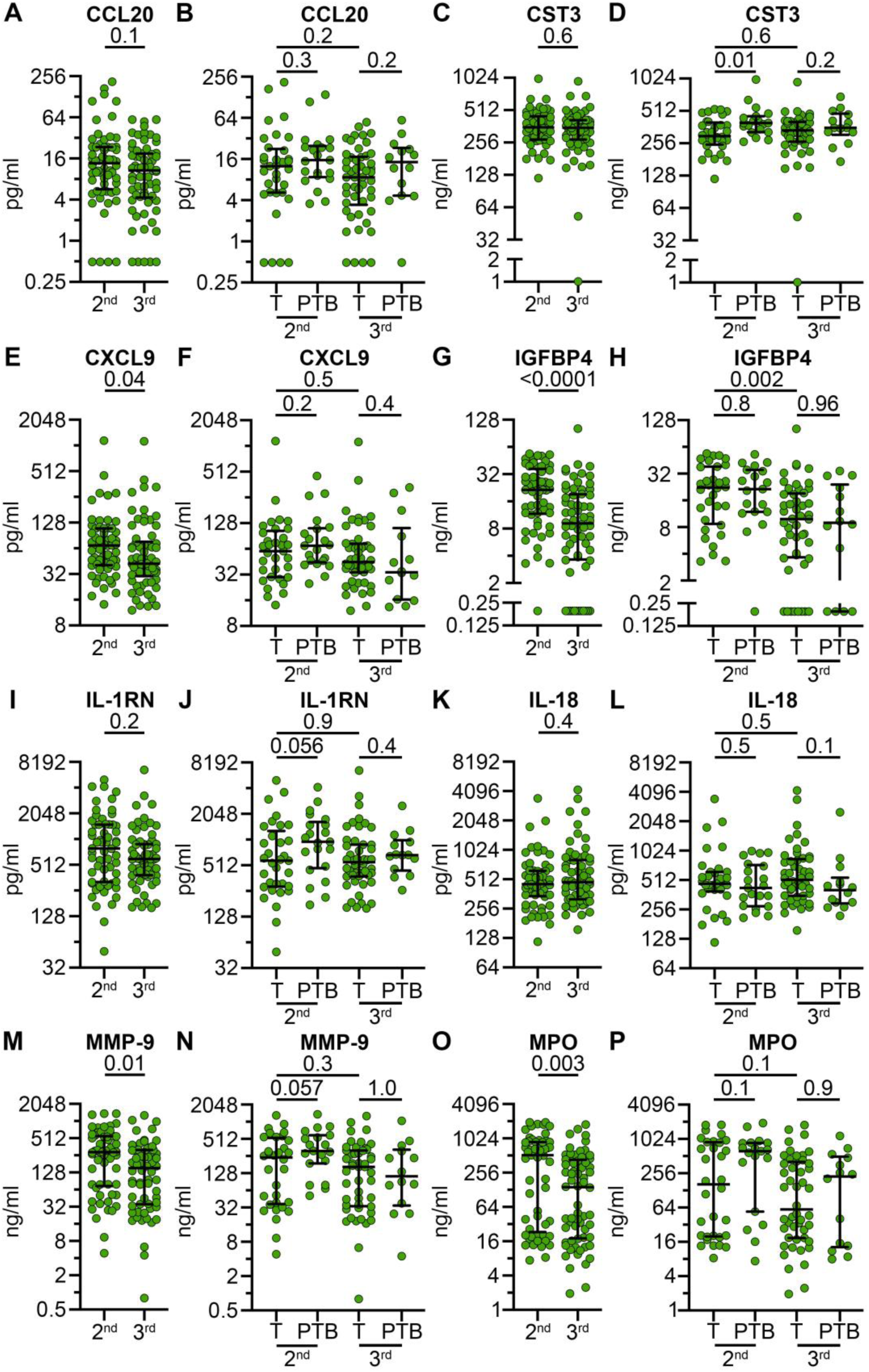
Trimester-specific immune factor concentrations in term versus preterm birth. Individuals were grouped by enrollment in second or third trimester, immune factors were assessed for significance by trimester, and within each trimester immune factors concentrations were compared between term and preterm birth. All immune factors that were significantly different by trimester, significantly (p<0.05) or non-significantly (0.05<p<0.1) associated with preterm birth at enrollment (Figure 2**, S8**) or significantly associated with preterm birth in one trimester or another were displayed (A, C, E, G, I, K, M, O). Individuals who gave birth at term (T) in the second (n=33) or third (n=50) trimesters were compared to individuals with a preterm birth (PTB) enrolled in the second (n=21) or third (n=14) trimesters (B, D, F, H, J, L, N, P). Statistical significance was tested by two-tailed Mann-Whitney (A, C, E, G, I, K, M, O) or Kruskal-Wallis uncorrected for multiple comparisons (B, D, F, H, J, L, N, P).

**Figure S9.**
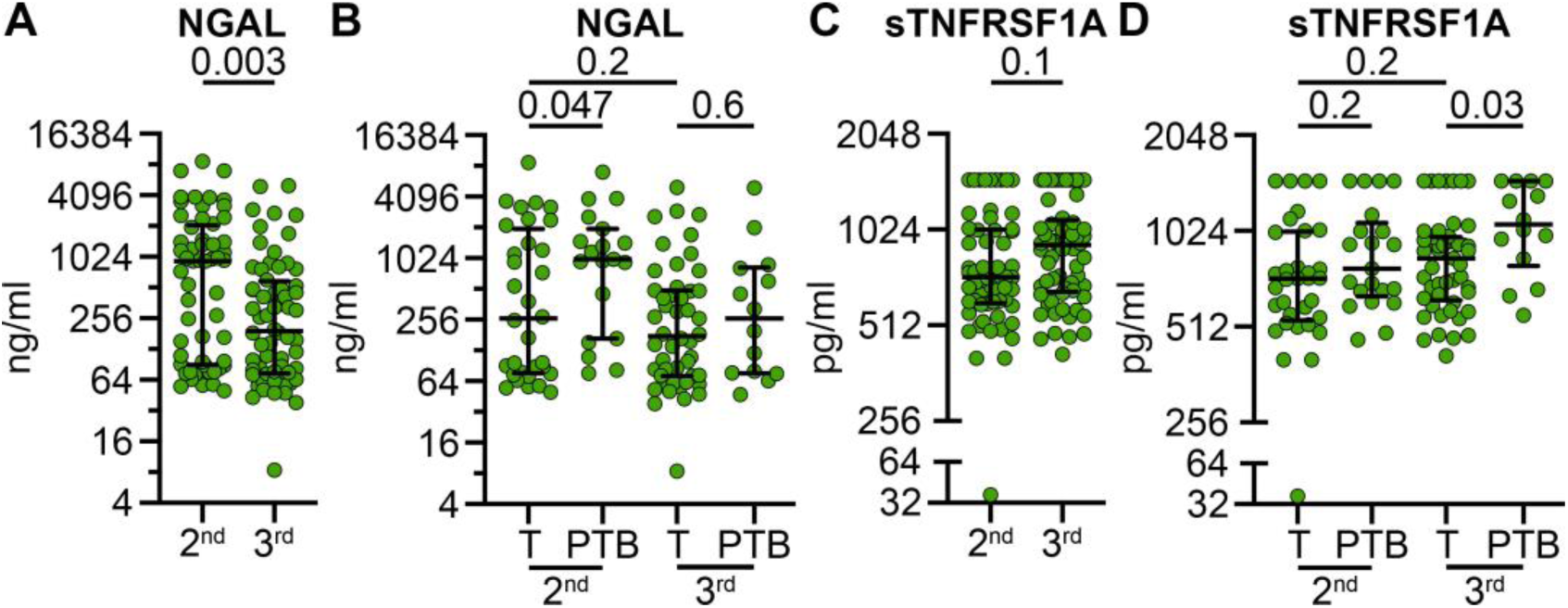
continued. Trimester-specific immune factor concentrations in term versus preterm birth. Individuals were grouped by enrollment in second or third trimester, immune factors were assessed for significance by trimester, and within each trimester immune factors concentrations were compared between term and preterm birth. All immune factors that were significantly different by trimester, significantly (p<0.05) or non-significantly (0.05<p<0.1) associated with preterm birth at enrollment (Figure 2**, S8**) or significantly associated with preterm birth in one trimester or another were displayed (A, C). Individuals who gave birth at term (T) in the second (n=33) or third (n=50) trimesters were compared to individuals with a preterm birth (PTB) enrolled in the second (n=21) or third (n=14) trimesters (B, D). Statistical significance was tested by two-tailed Mann-Whitney (A, C) or Kruskal-Wallis uncorrected for multiple comparisons (B, D).

**Figure S10.**
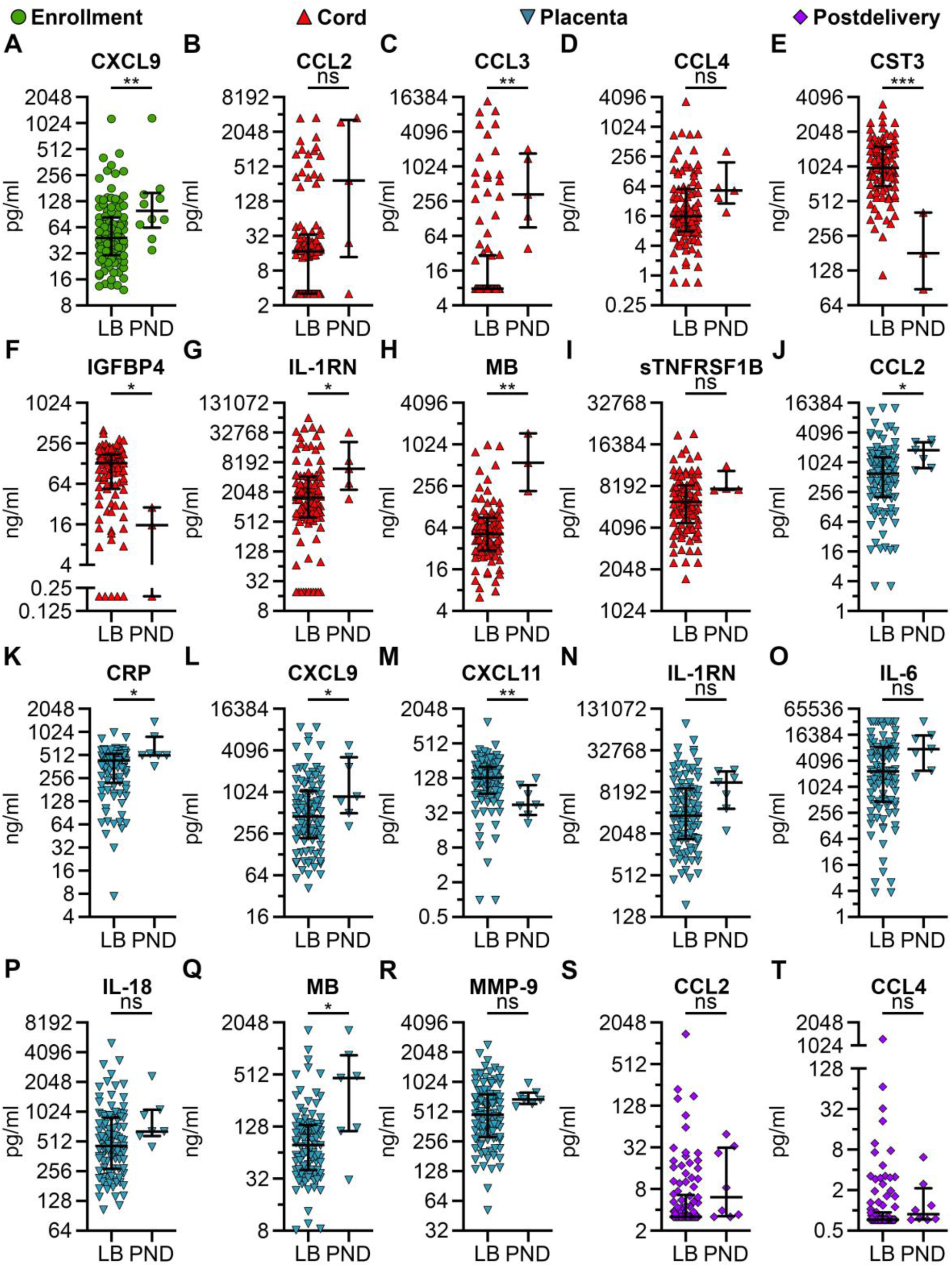
Immune factors associated with perinatal death. Immune factor concentrations were measured in plasma collected from participants at enrollment and postdelivery timepoints, as well as from placenta and cord blood. Concentrations were compared between live births (LB, n = 108) and perinatal deaths (PND, n = 10). Each point represents an individual participant’s analyte concentration; horizontal lines indicate the median and IQR. Statistical significance was assessed using a two-tailed Mann–Whitney test. *p<0.05, **p<0.01, ***p<0.001, ns signifies a p value between 0.05 and 0.1. See **Table S7** for descriptive statistics.

**Figure S10.**
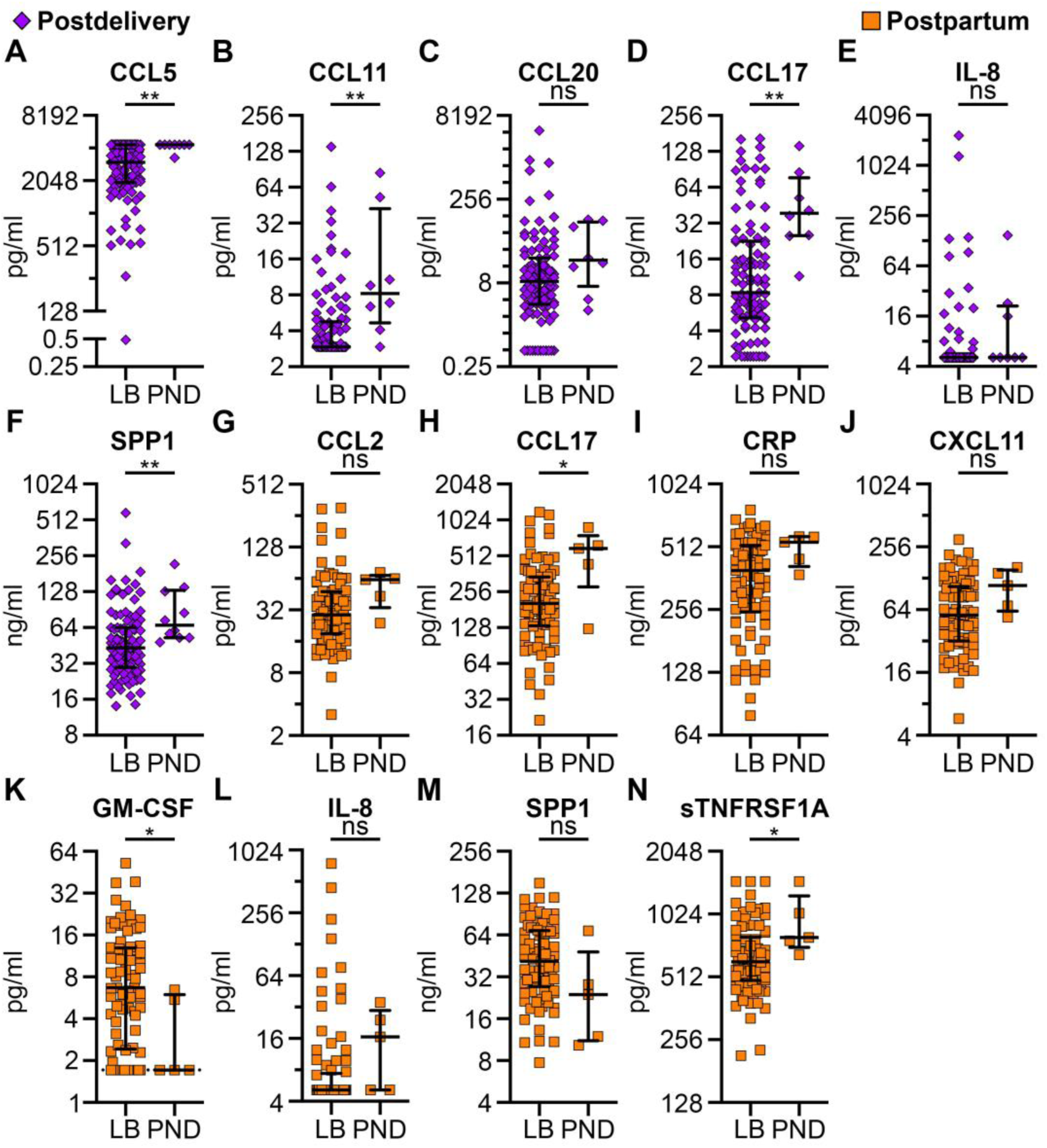
continued. Immune factors associated with perinatal death. Immune factor concentrations were measured in plasma collected from participants at postdelivery, and postpartum timepoints. Concentrations were compared between live births (LB, n = 108) and perinatal deaths (PND, n = 10). Each point represents an individual participant’s analyte concentration; horizontal lines indicate the median and IQR. Statistical significance was assessed using a two-tailed Mann–Whitney test. *p<0.05, **p<0.01, ns signifies a p value between 0.05 and 0.1. See **Table S7** for descriptive statistics.

**Figure S11.**
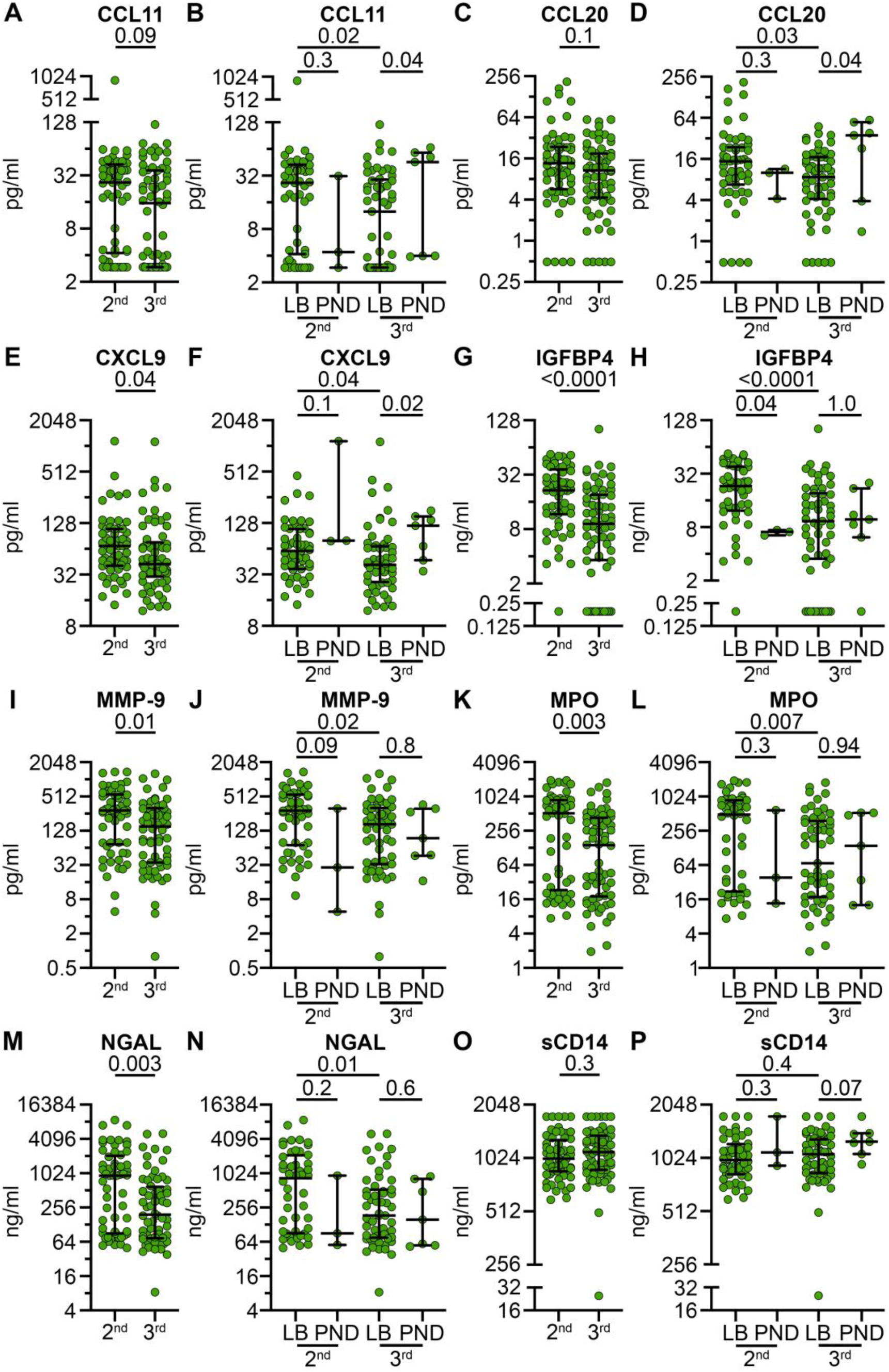
Trimester-specific immune factor concentrations in live birth versus perinatal death. Individuals were grouped by enrollment in second or third trimester, immune factors were assessed for significance by trimester, and within each trimester immune factors concentrations were compared between live birth and perinatal death. All immune factors that were significantly different by trimester, significantly (p<0.05) or non-significantly (0.05<p<0.1) associated with perinatal death at enrollment (Figure 3**, S10, S10 continued**) or significantly associated with perinatal death in one trimester or another were displayed (A, C, E, G, I, K, M, O). Individuals that had a live birth (LB) in the second (n=51) or third (n=57) trimesters were compared to those who had a stillbirth or neonatal death (PND) in the second (n=3) or third (n=7) trimesters (B, D, F, H, J, L, N, P). Statistical significance was tested by two-tailed Mann-Whitney (A, C, E, G, I, K, M, O) or Kruskal-Wallis uncorrected for multiple comparisons (B, D, F, H, J, L, N, P).

**Figure S12.**
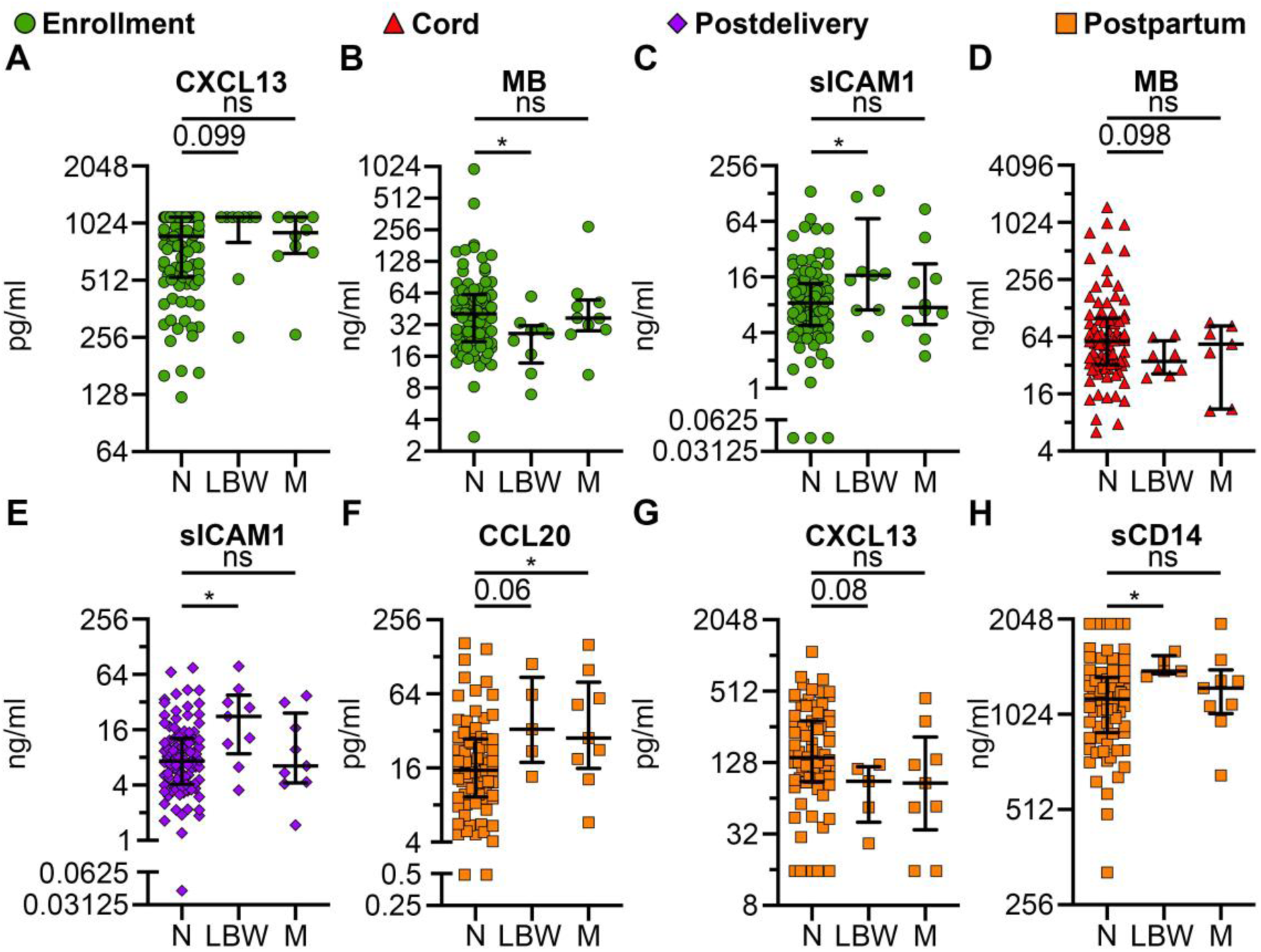
Immune factors associated with low birthweight. Immune factor concentrations were measured in plasma collected from participants at enrollment, postdelivery, and postpartum timepoints, as well as from cord blood. Concentrations of analytes were compared between individuals with normal birthweight (N, n = 99), low birthweight (LBW, n = 9), and those with missing birthweight data (M, n = 10). Each point represents an individual participant’s analyte concentration; horizontal lines indicate the median and IQR. Statistical significance tested by Kruskal-Wallis uncorrected for multiple comparison. *p<0.05, p-values between 0.05 and 0.1 labeled, ns=p-value>0.1. See **Table S8** for descriptive statistics.

**Figure S13.**
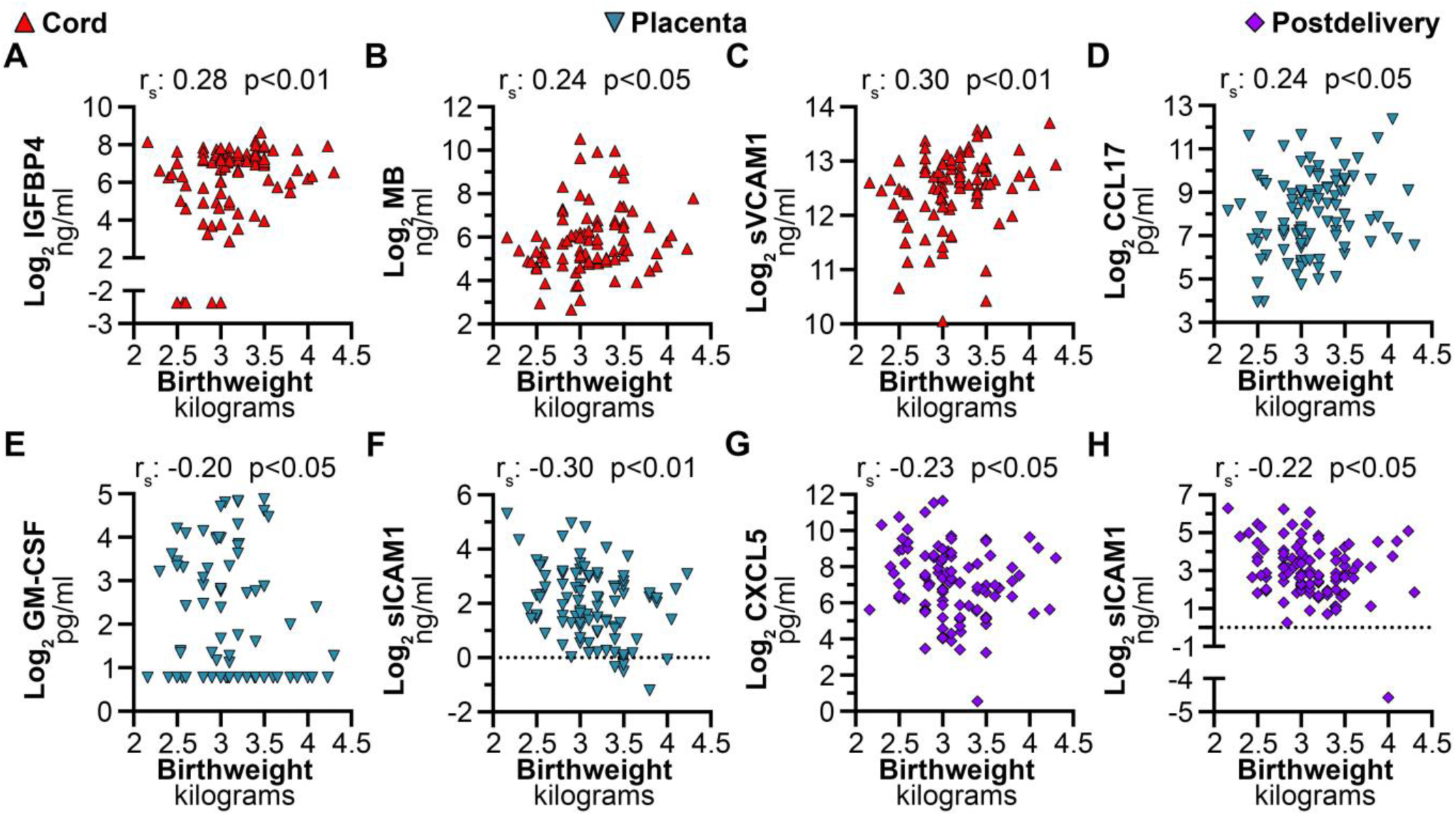
Immune factors that significantly correlate with birth weight. Log_2_ concentration of immune factors were plotted against the birthweight in kilograms. Each dot represents an individual participant’s analyte measurement. Statistical significance tested by Spearman correlation (r_S_) and approximate p-value of the correlation displayed. Only those with a p-value<0.05 and an r_S_>0.2 or <-0.2 were considered significant. Horizontal dotted lines represent the point at which y=0.

**Figure S14.**
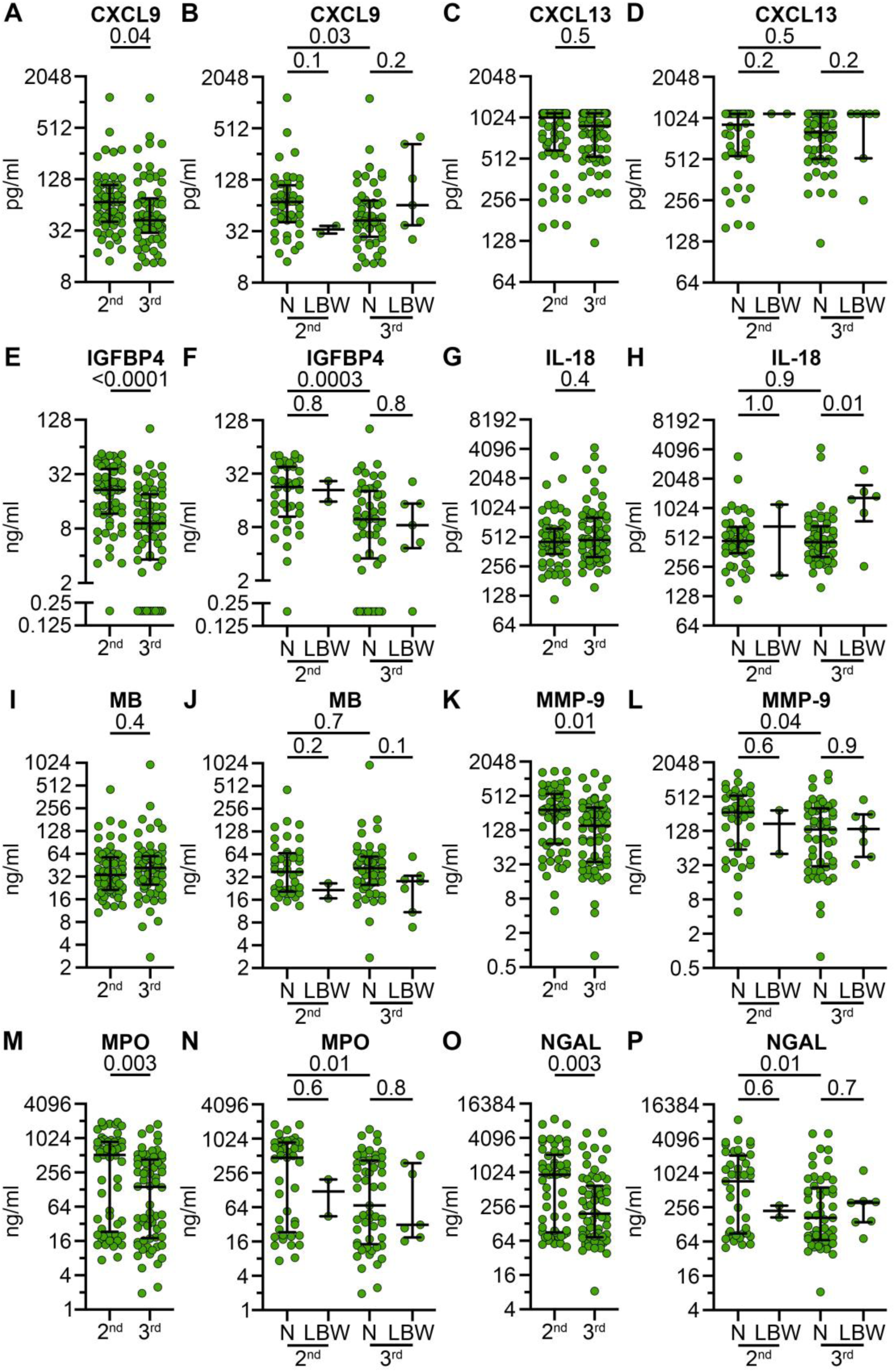
Trimester-specific immune factor concentrations in normal versus low birth weight. Individuals were grouped by enrollment in second or third trimester, immune factors were assessed for significance by trimester, and within each trimester immune factors concentrations were compared between normal (N) and low birth weight (LBW). All immune factors that were significantly different by trimester, significantly (p<0.05), non-significantly (0.05<p<0.1) associated with low birth weight at enrollment (Figure 4**, S12**) or significantly associated with low birth weight in one trimester or another were displayed (A, C, E, G, I, K, M, O). Individuals who gave birth to normal birth weight neonates in the second (n=44) or third (n=55) trimesters were compared to those who gave birth to low birth weight neonates in the second (n=2) or third (n=7) trimesters (B, D, F, H, J, L, N, P). Statistical significance was tested by two-tailed Mann-Whitney (A, C, E, G, I, K, M, O) or Kruskal-Wallis uncorrected for multiple comparisons (B, D, F, H, J, L, N, P).

**Figure S14.**
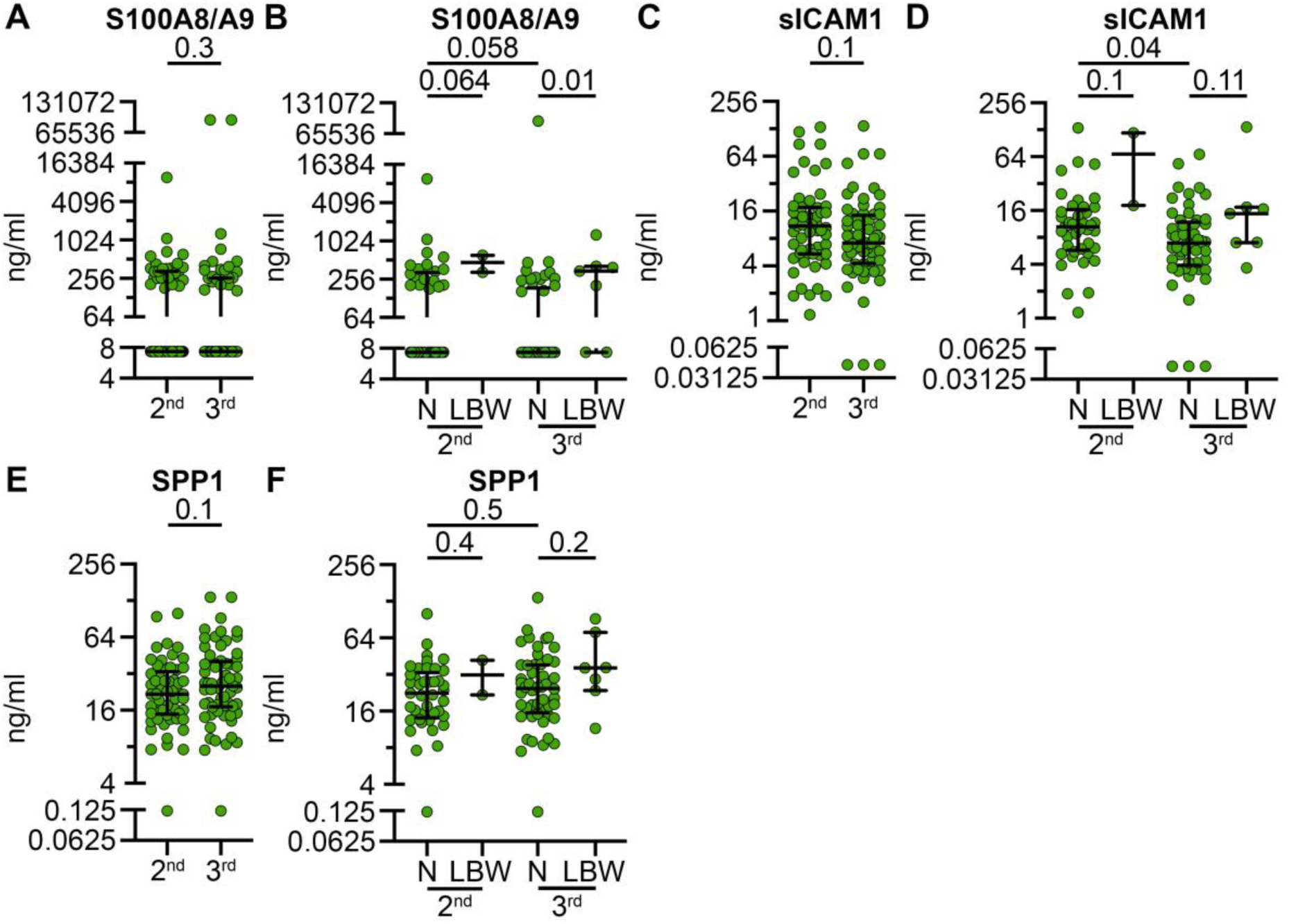
continued. Trimester-specific immune factor concentrations in normal versus low birth weight. Individuals were grouped by enrollment in second or third trimester, immune factors were assessed for significance by trimester, and within each trimester immune factors concentrations were compared between normal (N) and low birth weight (LBW). All immune factors that were significantly different by trimester, significantly (p<0.05) or non-significantly (0.05<p<0.1) associated with low birth weight at enrollment (Figure 4**, S12**) or significantly associated with low birth weight in one trimester or another were displayed (A, C, E). Individuals who gave birth to normal birth weight neonates in the second (n=44) or third (n=55) trimesters were compared to those who gave birth to low birthweight neonates in the second (n=2) or third (n=7) trimesters (B, D, F). Statistical significance was tested by two-tailed Mann-Whitney (A, C, E) or Kruskal-Wallis uncorrected for multiple comparisons (B, D, F).

**Figure S15.**
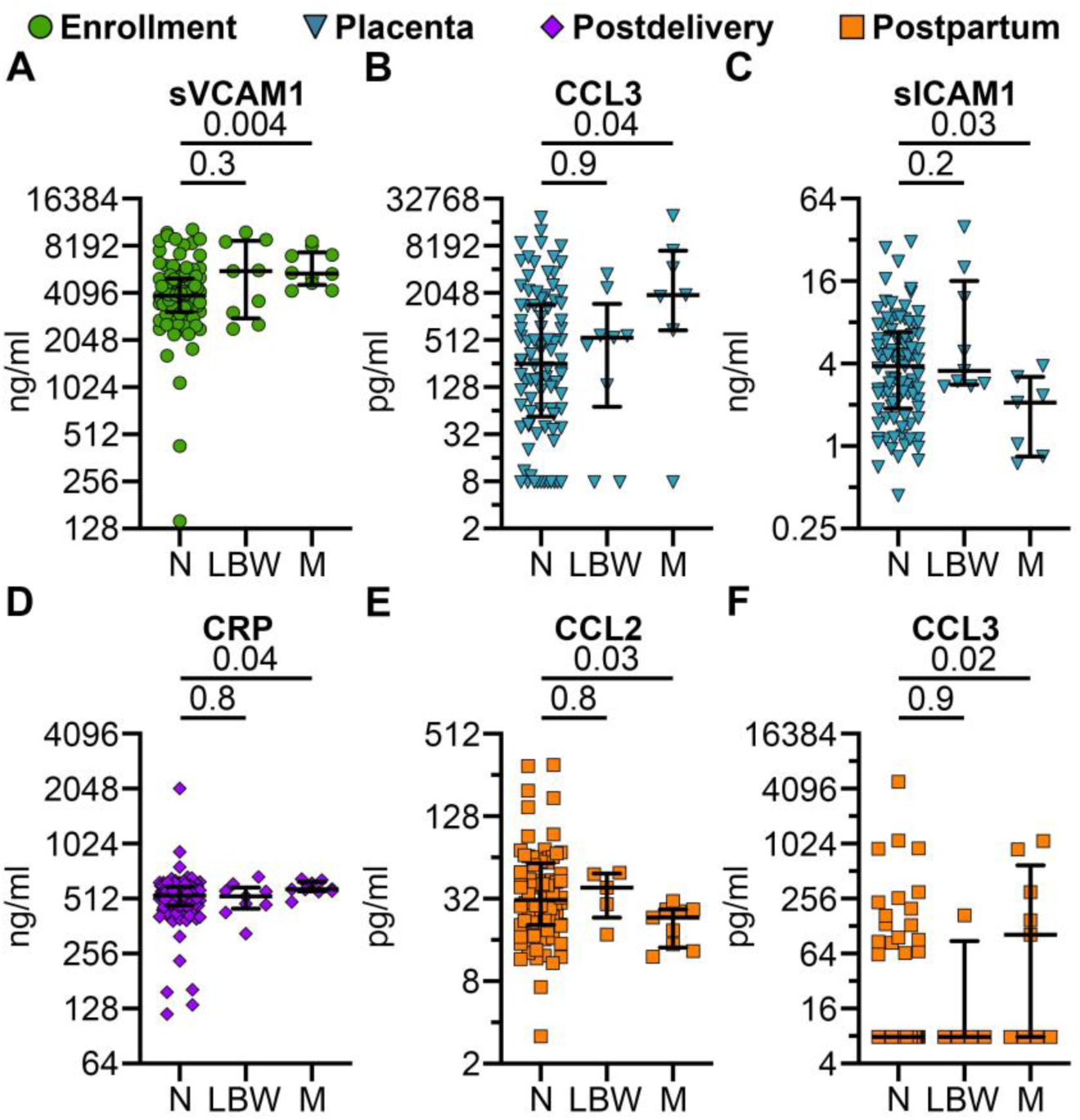
Immune factors associated with missing birth weights. Immune factor concentrations were measured in plasma collected from participants at enrollment, postdelivery, and postpartum timepoints, as well as from placental plasma. Concentrations of analytes were compared between individuals with normal birthweight (N, n = 99), low birthweight (LBW, n = 9), and those with missing birthweight data (M, n = 10). Each point represents an individual analyte concentration; horizontal lines indicate the median and IQR. Statistical significance tested by Kruskal-Wallis uncorrected for multiple comparison. Normal birth weight (N), low birthweight (LBW), missing birthweight (M).

**Figure S16.**
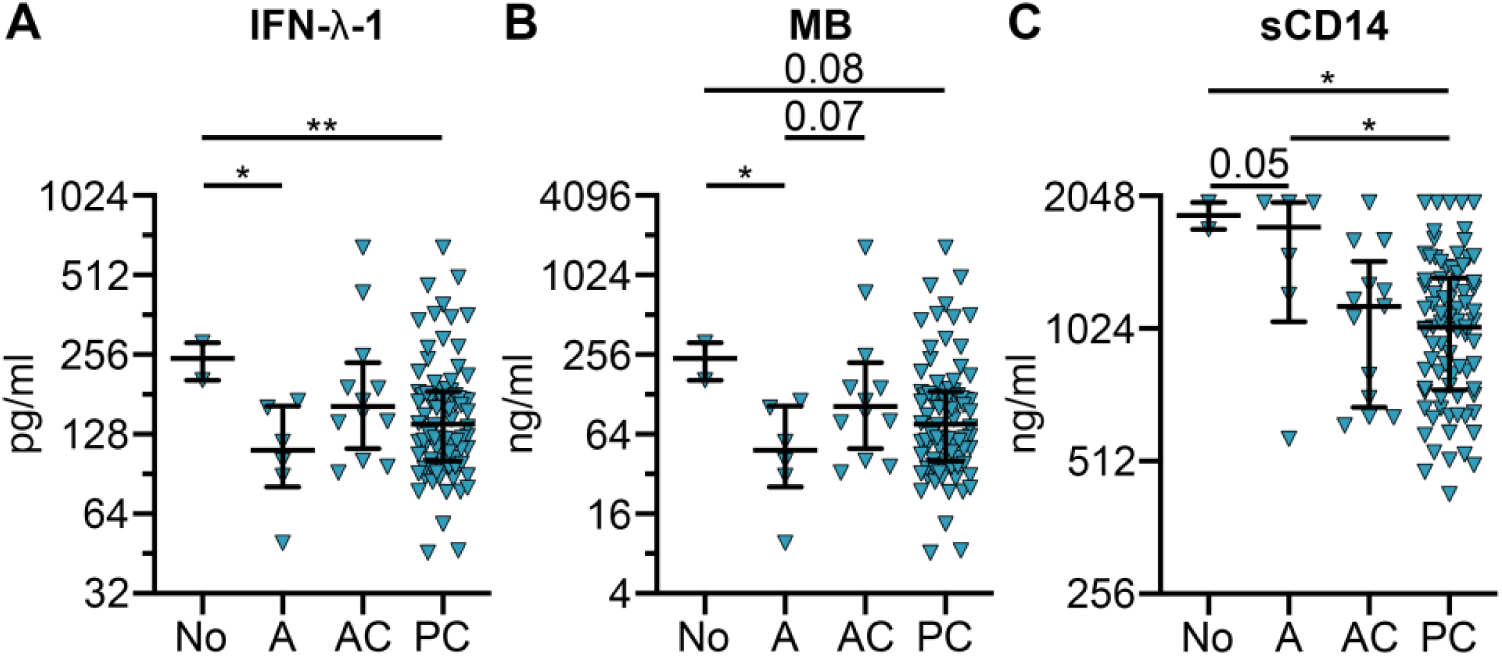
Comparison of immune factors based on placental malaria status. Levels of immune factors in placental plasma were analyzed across four groups defined by placental malaria status: none observed (No, n=2), active (A, n=6), active-chronic (AC, n=12), and past-chronic (PC, n=81). Three subjects lacking placental malaria data were excluded from this analysis. Significance was determined using a Kruskal-Wallis test, uncorrected for multiple comparisons. The figure displays only those immune factors demonstrating statistical significance or a p-value (0.05 < p < 0.1). Significance levels are denoted as *p < 0.05 and **p < 0.01.

